# Innate immune cell response to host-parasite interaction in a human intestinal tissue microphysiological system

**DOI:** 10.1101/2022.01.27.478058

**Authors:** Mouhita Humayun, Jose M Ayuso, Keon Young Park, Bruno Martorelli Di Genova, Melissa Skala, Sheena C Kerr, Laura J. Knoll, David J. Beebe

## Abstract

Protozoan parasites that infect humans are widespread, and lead to varied clinical manifestations, including life-threatening illnesses in immunocompromised individuals. Animal models have provided insight into innate immunity against parasitic infections, however, species-specific differences and complexity of innate immune responses make translation to humans challenging. Thus, there is a need for novel *in vitro* systems that can elucidate mechanisms of immune control and parasite dissemination. We have developed a human microphysiological system of intestinal tissue to evaluate parasite-immune-specific interactions during infection, which integrates primary intestinal epithelial cells and immune cells to investigate the role of innate immune cells during epithelial infection by the protozoan parasite, Toxoplasma gondii, which affects billions of people worldwide. Our data indicate that epithelial-infection by parasites stimulates a broad range of effector functions in neutrophils and NK cell-mediated cytokine production that play immunomodulatory roles, demonstrating the potential of our system for advancing the study of human-parasite interactions.

**Teaser:** Novel engineered model of human intestinal tissue for study of dissemination and immune control of parasitic infections.

## Introduction

Infections caused by parasitic pathogens are a global health problem that affects more than a quarter of the world’s population [1], yet effective antiparasitic therapeutics are limited and often come with severe adverse reactions. Similarly, vaccines are limited for any food or water-borne parasitic infection [2]. Immune responses initiated by the innate immune system in the intestinal tissue are a part of the frontline defense against parasitic infections. A more complex picture of their role in shaping protective immunity development and pathogenesis are beginning to emerge. Infection by the protozoan and intracellular parasite *Toxoplasma gondii* affects about a third of humans worldwide making it one of the most widespread human pathogens in the world [3]. Human exposure to *T. gondii* infection occurs primarily through the oral route due to ingestion of contaminated food or water [4]. However, there are significant knowledge gaps in how the innate immune system interacts with human intestinal parasites like *T. gondii* in the local microenvironment of the intestinal tissue that either reduce parasite dissemination or contribute to the development of invasive systemic disease. As responses to parasitic infection by the innate immune system play key roles in shaping protective immunity, improved knowledge of the mechanisms that initiate innate immunity and contribute to pathogenesis are central to developing effective therapeutics and vaccines.

The gut epithelium and vascular barrier regulate what enters the host tissue beyond the intestinal epithelial barrier and what enters the circulation. Functionally, the intestinal epithelial barrier separates the luminal contents from immune cells found in the gut parenchyma and prevents the systemic dissemination of the microbiota and enteric pathogens to liver, spleen and other peripheral tissues [5]. During parasitic infection, immune cells in the gut-parenchyma coordinate host-protective responses necessary for resolving acute infection and preventing tissue-dissemination [6]. Currently, murine hosts are predominantly used as translational models for *T. gondii* infections in humans. Despite the vast knowledge of basic and translational human immunology obtained from mouse studies, several components of the mouse immune system are incongruent with the human immune system [7,8]. Humanized mice allow for improved modelling of the human immune systems and have emerged as alternatives to traditional rodent models. However, challenges still remain due to the lack of critical adhesion molecules and other signaling proteins in humanized mouse models that are required for mounting human-specific immune responses to infections [9,10]. While *in vivo* animal models replicate microenvironmental complexity and physiological conditions, several aspects including species-specific differences, limited ability to image pathogen trafficking and the time/expense of mechanistic study makes investigation of pathogen trafficking challenging in these models. Thus, there is a need for relevant human *in vitro* models to interrogate the innate immune responses to pathogens.

Organ-on-chip devices and microphysiological systems (MPSs) have overcome some of the challenges of traditional *in vitro* approaches by enabling the integration of three-dimensional (3D) complexity, spatial organization and relevant cell-cell and cell-extracellular matrix interactions for modeling *in vivo-*like responses [11–13]. Several technologies for modeling pathophysiological processes in human tissue have emerged that represent important steps forward in the evolution of organ-on-a-chip platforms. MPS of the lung [14–16], brain [17], bladder [18], liver [19] and intestine [20–25] have been developed for studies of host interactions with viruses, fungi, bacteria, and parasites [26,27]. In the context of modeling host-microorganism interactions, viral infections and bacterial colonization in the epithelia have been the primary applications of MPSs, with emerging applications in testing antiviral therapeutics and modeling innate immune responses. Immune cell trafficking across endothelial and epithelial barriers have been studied using these models [22,28], but difficulties persist with integrating membrane-free interfaces that mimic tissue composition of the gut epithelial and vascular barrier, including cellular and extracellular matrix (ECM) components, and tissue architecture (i.e., endothelial vessel, epithelial lumen geometry) that can critically influence host and pathogen responses. Recent efforts to model the 3D tissue anatomy of intestinal epithelia have culminated in the establishment of organoid-on-a-chip systems that retain the cellular diversity and regenerative potential offered by organoid technology while addressing the problem of lumen accessibility using microengineering or tissue engineering approaches [29–31]. Using advanced 3D bioprinting techniques, culture of intestinal tubes can be maintained long-term in ECM gels within these systems [29]. These systems can support *in vitro* tissue homeostasis of the intestinal epithelium, and its interactions with underlying tissue components. However, functional responses of innate immune cells and concomitant molecular analysis of responses to microorganisms in these models have not been characterized.

In this work, we developed a novel MPS of the human intestinal tissue to study host-parasite interactions and innate immune cell response to parasite infection *in vitro*. By using a highly tractable micromolding technique for creating hollow structures, we generated tubes of intestinal epithelium and endothelium supported by ECM gel, recapitulating the lumen geometries of the gastrointestinal tract and blood vessels. We microengineered a human-relevant model of the gut-epithelium and vascular barrier, within the intestinal tissue, that provides means for studying gastrointestinal parasitic infections and the associated interactions with innate immune cells. We demonstrate adaptation of this model to incorporate primary human intestinal stem cells from organoids derived from intestinal tissue resections. We used the system to model parasite invasion and replication within the intestinal epithelium, followed by leukocyte trafficking to the site of infection. Neutrophils and NK cells are critical for early protection and reducing parasite burden during *T. gondii* infection[32,33]. By measuring gene transcription, metabolic changes, and cytokine secretions, we gained interesting insights into the function of human innate immune cells, including neutrophils and NK cells, during the initial stages of *T. gondii* parasite infection of the intestinal epithelium. Hence, the engineered MPS model of the intestinal tissue emulates critical features of host-parasite interactions enabling us to address questions related to human-relevant tissue response, activation of innate immunity, and the role of immune cells in parasite dissemination.

## Results

### Development of an intestinal tissue MPS for studying host-pathogen interactions

Infection by *T. gondii* is naturally acquired through oral ingestion of food or water that are infected with parasite cysts or oocysts [34]. The intestinal epithelium, characterized by a villus-crypt axis consisting of a single layer of constantly renewing and differentiating epithelial cells, separates microbes in the lumen from the intestinal vascular systems, [35,36] **Figure 1A**. Experiments in mice and cell line/explant studies have shown that translocations of *T. gondii* from the apical surface of the intestinal epithelium to the basal side, during acute infection, may occur via epithelial transmigration or following epithelial invasion and intracellular replication [37,38]. Tissue damage caused by the *T. gondii* epithelial invasion and intracellular replication leads to the secretion of a host of inflammatory factors by intestinal tissue-resident cells [39], **Figure 1A**. Presence of these factors increases the expression of integrins and chemokine receptors on endothelial cells of local vasculature that promote immune cell extravasation into the lamina propria. Leukocytes, primarily neutrophils, are then recruited to the site of infection from neighboring vasculature where they encounter effector-enhancing cytokines and pathogen-derived products to neutralize the parasite by killing or controlling their replication [34,40–42], **Figure 1A**. Other innate immune cells such as NK cells are also recruited from the vascular system that further amplifies the inflammatory response by producing effector-enhancing cytokines[43,44], **Figure 1A**. A robust immune response during these early stages of the *T. gondii* infection is critical for shaping innate immunity which in turn induces acquired immunity.

**Figure 1.**
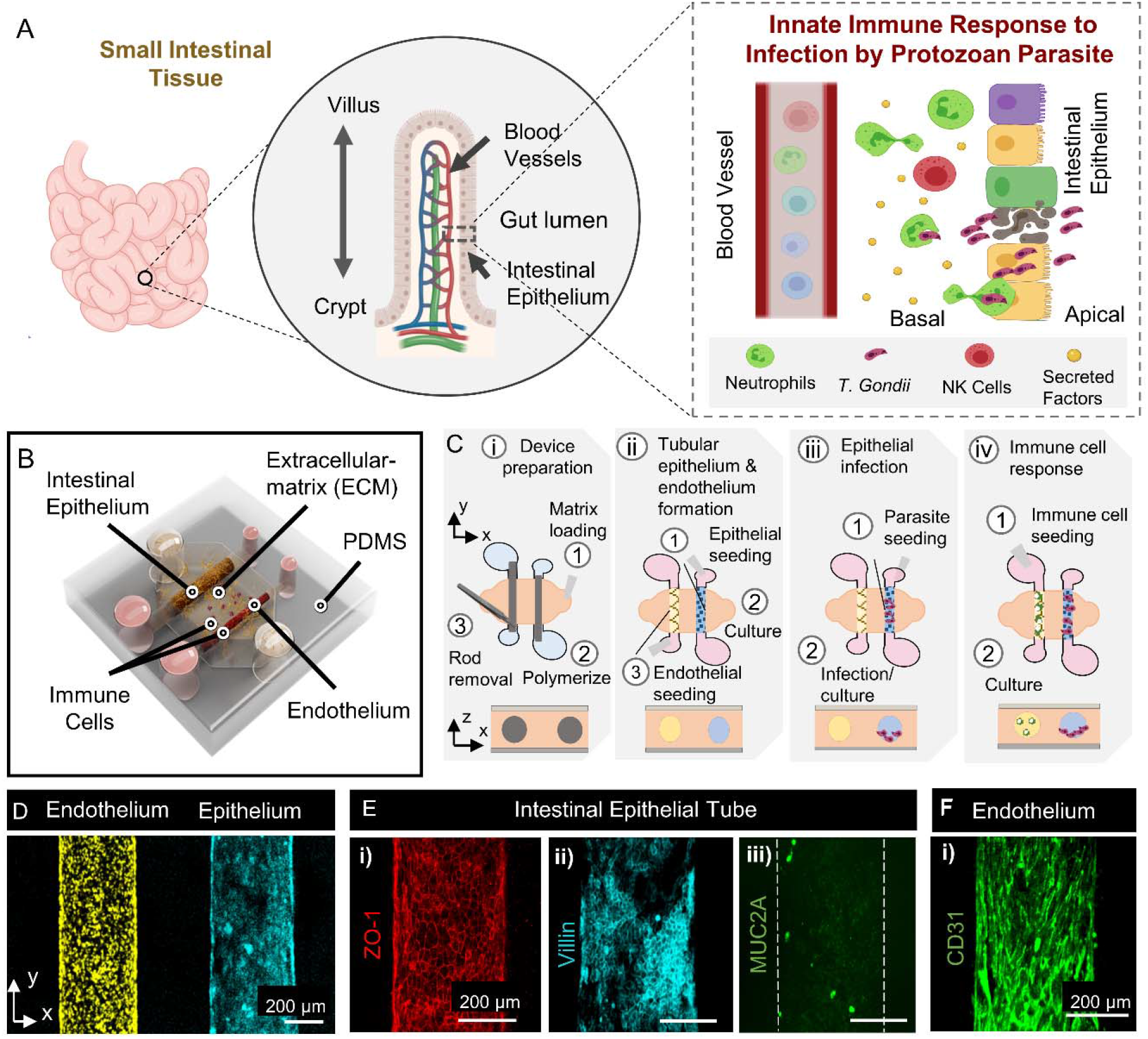
Human intestinal tissue microphysiological system for studying innate immune responses to parasitic infection. **(A)** Schematic representation of the design rationale for modeling parasite infection of the intestinal epithelium and innate immune cell responses. **(B)** 3D rendered illustration of the intestinal tissue MPS which include tubular intestinal epithelium and endothelium within an ECM gel. Immune cells are introduced into the lumen of the endothelium for modeling and elucidating innate immune responses to parasite infection of the epithelium. **(C)** Schematic representation describing the experimental approach to setup each component (i-iv) of the MPS used in this study. **(D)** Fluorescence image showing the formation of a confluent Caco-2 intestinal epithelium (yellow) and HUVEC endothelium (cyan). **(E)** The model retains phenotypic characteristics of tight junction markers (i, ZO-1), microvilli markers (ii, villin) and markers of mucus producing goblet cells (iii, MUC 2A) for the epithelium. **(F)** HUVEC endothelium in the co-culture with the epithelium retains expression of endothelial marker (CD31).

To model these initial stages of *T. gondii* infection, we bioengineered an intestinal tissue MPS that incorporates 3D tubular structures of the intestinal epithelium and the endothelium, and immune cell components to allow the analysis of host immune responses against intestinal pathogens in a more physiologically relevant culture microenvironment, **Figure 1B**. Inspired by previous work by ourselves and others, we generated lumen structures within a gas and nutrient-permeable collagen-based ECM hydrogel using a micromolding technique for modeling tubular geometries of the intestinal epithelium and the gut vascular barrier within intestinal tissue[29,45– 48]. The hydrogels are integrated into a polydimethylsiloxane (PDMS) elastomeric device with a central chamber, that contains the ECM gel, connected to two pairs of inlet and outlet ports that support PDMS rods used for molding the tubular structures, **Figure 1C i**. Following ECM gel polymerization, the rods are pulled out of the chamber leaving behind hollow lumen structures. The inlet and outlet ports connected to the chamber provide direct access to the lumens which can be used for cell seeding and for the supply of medium and growth factors during culture. We injected and cultured intestinal epithelial cells inside one of the tubular structures to form an epithelial lumen with direct access to the apical surface. After the epithelial lumen is formed, in the same manner, we seeded and cultured vascular endothelial cells in the adjacent lumen to generate a biomimetic endothelial vessel, **Figure 1C ii**. To model epithelial invasion of *T. gondii*, parasites are seeded into the apical surface of the epithelium through the inlet port and cultured, **Figure 1C iii**. Next, immune cells are introduced directly into the endothelial vessel and cultured for evaluating the immune response, **Figure 1C iv**. Altogether, our model mimics critical components of the intestinal tissue including the epithelium, adjacent blood vessels, and immune cell components for studying responses to infection by protozoan parasites. For initial characterization of the model, we generated tubular intestinal epithelium using the human colon epithelial cell line, Caco-2 and formed a tube-shaped endothelial vessel using human umbilical vein endothelial cells (HUVECs), **Figure 1D**. The ECM hydrogel provides structural support for the formation of continuous layers of epithelial and endothelial cells that functions as a barrier between the apical side of the lumen and the underlying matrix. The intestinal epithelium generated within our system expresses tight junction marker ZO-1, which is involved in barrier function, as well as villin-1 which is involved in microvilli formation and epithelial restitution after damage, **Figure 1E i-ii**. The intestinal epithelium is increasingly recognized as a critical component of mucosal innate immunity against invading microorganisms through the secretion of mucin and antimicrobial proteins. We demonstrate the expression of mucin 2 (MUC2), which is secreted by goblet cells in the intestinal epithelium and forms a major component of the inner mucus layer, **Figure 1E iii**. Expression of these cell lineage markers indicates differentiation and maturation of the epithelium cultured within our system. To assess characteristic features of the endothelial vasculature, we stained for and observed the expression of endothelial junction protein CD31, **Figure 1F**. Together, these results show that co-cultures of tubular intestinal epithelium and endothelium supported by collagen-based ECM gel retain the phenotypic characteristics of a mature epithelium and endothelium.

### Generation and characterization of primary small intestinal tubes in the intestinal tissue model MPS

*In vitro* cultures of patient or iPSC-derived intestinal epithelial stem cells in 3D organoids can generate a fully differentiated and polarized epithelium [49–51]. However, the relative inaccessibility of the apical surfaces makes the inoculation of larger pathogens, such as *T. gondii* compared to bacteria or viruses, into the cavity difficult to perform. Moreover, the variable 3D geometry of organoids makes real-time imaging of host-pathogen interactions technically challenging. Therefore, we adapted our intestinal tissue MPS to integrate the formation of tubular epithelium using primary human intestinal epithelial cells, from intestinal organoid fragments, to provide a model that retains the major epithelial cell types and provides easy access to the apical surface, **Figure 2A**. To incorporate primary human intestinal epithelial cells, we obtained surgical samples from macroscopically normal regions of the human small intestinal tissue. Human intestinal crypts containing functional stem cells, derived from the jejunum region of the small intestine, were used to generate organoids, and cultured for > 5 passages prior to use within our model. Jejunal organoids cultured in an expansion medium for 7-9 days were dissociated into a mix of fragments and single-cell suspensions before being introduced into the lumen tube of the microdevice, **Figure 2B**. To maximize coverage of the luminal surface, primary small intestinal cells were seeded into the device twice and cultured in expansion media for 24h between each cell seeding. The device was flipped upside down prior to the second round of cell seeding to facilitate cell adhesion to the top half of the lumen surface. Initially, the large majority of cells appear spread across the luminal surface within 1-2 days of cell seeding and progressively grow to form a continuous epithelium. Following cell adhesion to the matrix (approximately 72h from initial cell seeding), the culture medium is changed to a differentiation medium and the intestinal epithelium is cultured for 5-7 days to promote differentiation of epithelial subtypes. We utilized a previously published differentiation culture medium formulation and made further modifications **(see Materials and Methods)** to accommodate the culture of HUVECs in the adjacent lumen following differentiation of the epithelium. HUVECs were seeded into the adjacent lumen and the device was inverted every 30 min over a two-hour period to coat the lumen with endothelial cells. The tubes of intestinal epithelium and endothelium were co-cultured for an additional 48h prior to infection with *T. gondii*. For modeling immune response to parasite infection, the intestinal epithelial tube was infected with *T. gondii* for 48h and immune cells including neutrophils and NK cells were added to the lumen of the endothelium followed by co-culture within the model for 24-48h, **Figure 2B**.

**Figure 2.**
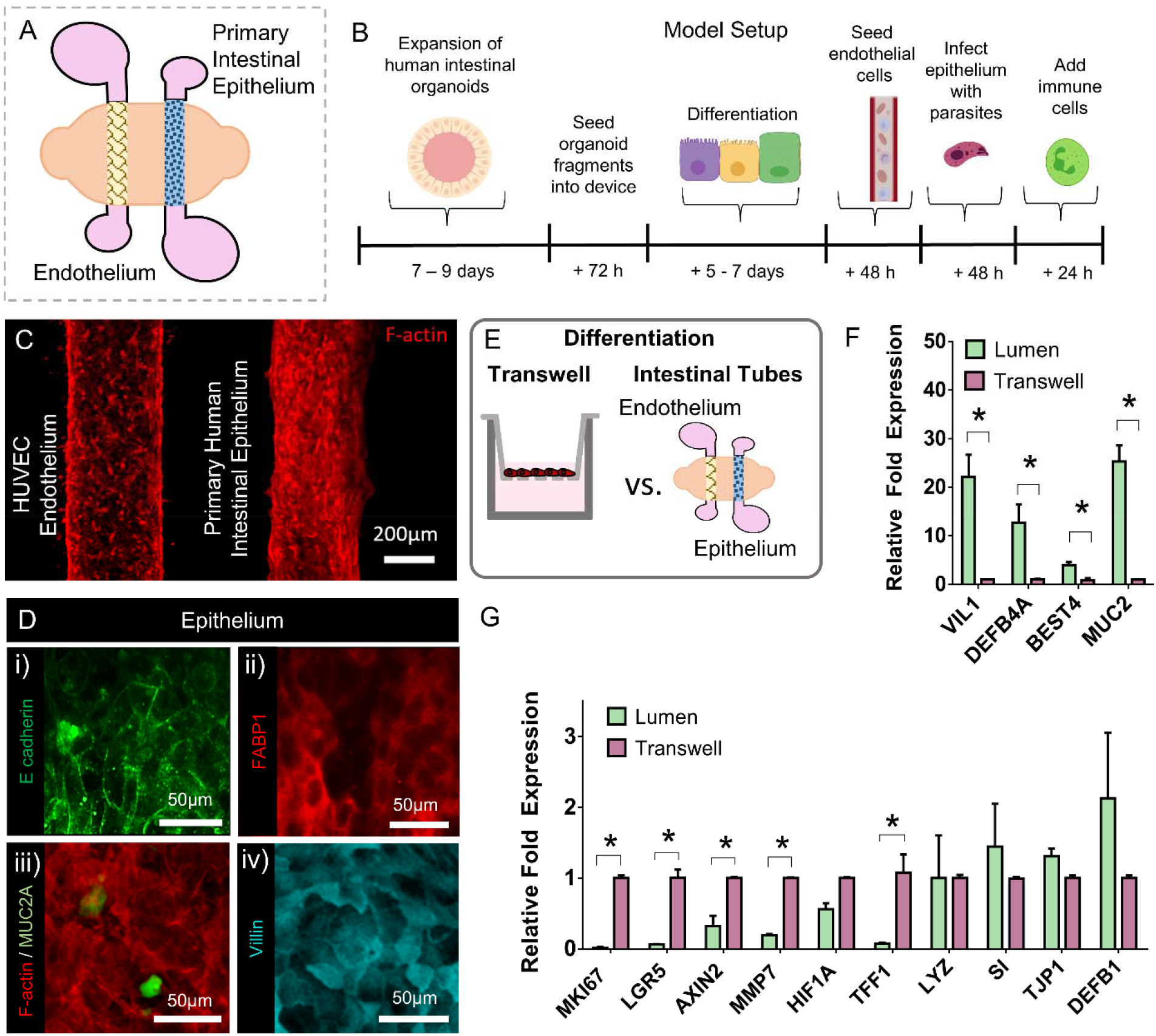
Integrating primary intestinal epithelial cells in the human intestinal tissue MPS. **(A)** Schematic representing the spatial distribution of the tubular intestinal epithelium and endothelium used in the intestinal tissue MPS. **(B)** Optimized culture protocol and timeline to setup the intestinal tissue MPS. This includes expansion of intestinal organoids, formation of intestinal tubes, differentiation of the epithelium, formation of the tubular endothelium, parasite infection of the epithelium and addition of immune cells into endothelium. **(C)** HUVEC endothelial cells and primary intestinal epithelial cells formed lumen structures in the intestinal tissue model as shown by immunostaining of F-actin (red). **(D)** Following culture and differentiation of the tubular epithelium, the intestinal epithelial cells show retention of phenotypic characteristics including expression of (i) epithelial cell-adhesion protein (E-cadherin, green), (ii) enterocyte-specific marker (FABP1, red), (iii) marker for mucin-producing cells (MUC 2A, green), and (iv) marker for protein involved in formation of microvilli (Villin 1, cyan). **(E)** Schematic representing the culture and differentiation of human primary intestinal epithelial cells in transwells and in the intestinal tissue MPS. **(F)** Bar graph showing differential gene expression of markers associated with proliferation and differentiation, and a functional intestinal epithelium in the intestinal tissue MPS versus in transwell culture. Genes analyzed include villin-1 (VIL1) for microvilli formation; β-defensin-4 (DEFB4) for antimicrobial peptides; Bestrophin-4 (BEST4) for absorptive cells and mucin 2 (MUC2) for goblet cells. **(G)** Bar graph showing gene expression of downregulated genes along with other markers of intestinal epithelial cells in the intestinal tissue model versus in transwell culture. Genes analyzed include proliferation marker (MKI67) for stem cells, leucine-rich-repeat-containing G-protein-coupled receptor 5 (LGR5) for intestinal stem cells; Axis Inhibition Protein 2 (AXIN2) for crypt regeneration; Matrix Metallopeptidase-7 (MMP7) bactericidal and anti-inflammatory effects; hypoxia-inducible factor-1 (HIF1A) for hypoxia; Trefoil Factor 1 (TIFF1) for mucosal repair; lysozyme (LYZ) for Paneth cells; sucrase-isomaltase (SI) specific for absorptive enterocytes; tight junction protein-1 (TJP1) for tight junctions in the epithelium and β-defensin-1 (DEFB1) for antimicrobial peptides. Values are presented as mean ±SD from 4 independent experiments involving tubular or monolayer epithelium generated from human intestinal organoids (asterisk denotes P value of ≤ □ 0.05).

The incorporation of human primary intestinal epithelial cells within our intestinal tissue model led to well-developed tubes of small intestinal epithelium and an adjacent vascular lumen, as indicated by immunofluorescent staining of F-actin, **Figure 2C**. Furthermore, the small intestinal epithelial lumen expressed E-cadherin and FABP1 which are typical markers of intestinal epithelial origin [52,53], **Figure 2D i-ii**. Similar to the tubes generated from the intestinal epithelial cell line, Caco-2, epithelial tubes generated primary intestinal epithelial cells exhibited positive expressions of MUC2 and villin, **Figure 2D iii-iv**.

*In vivo*, the intestinal epithelium rests on a supporting basement membrane composed of structural and adhesive proteins that epithelial cells use to anchor, migrate, and differentiate [54]. Matrix properties such as ECM ligands, stiffness and porosity are key factors that influence a wide range of cell behaviors including viability, tissue organization/architecture, and stem cell renewal and differentiation [55–58]. To further characterize the degree of differentiation induced in the intestinal tissue MPS, we conducted gene expression analysis of the intestinal tubes and compared the differentiation potential against epithelial monolayers grown in a standard culture platform (transwell), **Figure 2E**. Small intestinal epithelial cells in the intestinal tissue MPS and the transwell system were cultured under the same conditions: in the presence of expansion media for 2 days, followed by 7 days of culture in the differentiation medium. Reverse transcription and quantitative PCR (RT–qPCR) demonstrated that both the tubular (microphysiological model) and the monolayer (transwell) epithelium expressed region-specific marker, brush border enzymes such as sucrase-isomaltase (SI), which is native to jejunal tissue. SI gene expression was preserved beyond passage 25, indicating that the regional identities of organoids are intrinsically programmed. Genes associated with normal proliferation and differentiation, or functional human intestinal epithelial cells were compared, **Fig. S1-2**, and several genes were differentially expressed. RT-qPCR analysis revealed that jejunal epithelial differentiation in the intestinal tissue device resulted in increased expression of differentiation genes (VIL1 (marker for microvillar actin filament and enterocytic epithelial maturation), MUC2 (marker for mucin-producing cells), DEFB1 (marker for production of antimicrobial peptide, β-defensin 1), and SI) compared to differentiation in the transwell system, **Figure 2F**. Conversely, lower expression of proliferative genes (LGR5 (leucine-rich repeat-containing G-protein-coupled receptor 5, marker for stem cell renewal), KI67(a marker for proliferation), AXIN 2 (a marker for crypt regeneration), and MMP7(a known marker of Paneth cells, required for activating cell defensins [59]) were found in the intestinal tissue model relative to the transwell system,[59] **Figure 2G**. These results show that small intestinal epithelial cultures in the intestinal tissue model display features of their intestinal regional identity, and compared to standard culture techniques, exhibit improved differentiation characteristics/potential, and attenuated proliferative characteristics.

### Modeling protozoan parasite infection in human intestinal epithelium

Having established that the intestinal tissue MPS recapitulates tubular geometries of the intestinal epithelium and vasculature, the system has strong potential for modeling parasite invasion, replication, and translocation beyond epithelium into the lamina propria, **Figure 3A**. *T. gondii* infects the small intestinal epithelium cells following ingestion of parasites found in contaminated meat, produce, or water [34]. During acute infection, *T. gondii* parasites that invade the intestinal epithelium differentiate into a rapidly replicating life-stage, referred to as the tachyzoite stage, which then disseminates throughout the body [34,60]. To model *T. gondii* infection of the intestinal epithelium, we introduced tachyzoites into the tubular epithelium of our intestinal tissue MPS. Our model provides an exposed luminal surface accessible to pathogens, allowing us to directly expose the cells to the parasite. Caco-2 epithelial tubes were exposed to approximately 8 × 10^7^ transgenic mCherry-tagged *T. gondii (*type II ME49 strain) tachyzoites per tube. The lumen of the epithelial tube was washed with media 16h post-infection (hpi) to remove parasites that did not adhere to or invade the epithelium. Discrete and dense areas of parasites could be observed within a small proportion of epithelial cells using fluorescence microscopy at 24hpi and were largely contained within the boundaries of the epithelial tube, **Figure 3B**. The proportion of epithelial cells harboring the parasites increased between 24hpi and 48hpi and epithelial cell lysis was observed at 72hpi, **Figure 3C**, releasing tachyzoites into the lumen of the epithelium and the basal side of the epithelium. To confirm and quantify parasite replication, SAG1 (*T. gondii* surface antigen) gene expression levels were assessed at 24 hours and 48 hours via qPCR, **Figure 3D**. SAG1 expression analysis in the infected epithelium showed a 6-fold increase at 48hpi relative to 24hpi. Together, these results demonstrate that tubular intestinal epithelium generated in our human intestinal tissue MPS can support the invasion, replication, and translocation of *T. gondii* beyond the epithelium, which are key initial stages of the infection process prior to the encounter with innate immune cells.

**Figure 3.**
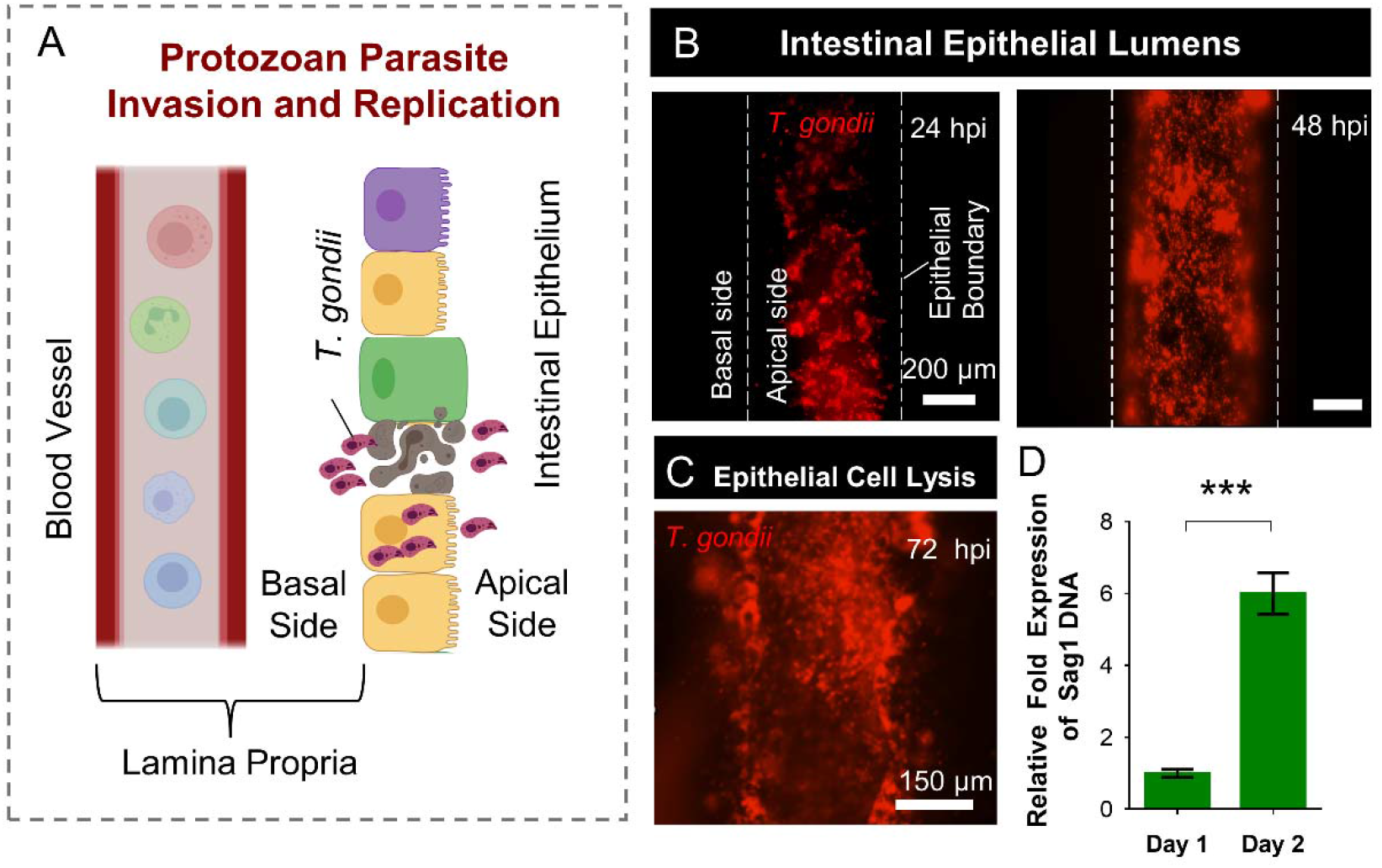
Modeling protozoan parasite invasion and replication in the intestinal epithelium. **(A)** Schematic representation of epithelial infection by *T. gondii*. Infection involves epithelial invasion, intracellular replication and transmigration or epithelial cell lysis of *T. gondii*, which release the parasites into the lamina propria containing immune cells and intestinal vasculature. **(B)** Caco-2 epithelial tubes were infected with m-cherry tagged *T*.*gondii* of the ME49 strain for 24h and imaged by fluorescence microscopy. The images depict time course images at 24h (i) and 48h(ii), showing that viable *T. gondii* were present within the epithelial lumen. White dotted line represents the epithelial boundary separating apical (lumenal) surface from the basal surface. **(C)** Gene expression analysis of genomic DNA from the epithelium for tachyzoite marker SAG1 shows significant up-regulation (approximately 6-fold expression) after 48h of infection relative to 24h. Values are presented as mean ±SD from 2 independent experiments and 12 different devices (asterisk denotes P value of ≤ □0.001). **(D)** Fluorescent images depicting epithelial cell lysis 72h following infection with m-cherry tagged *T*.*gondii*.

### Neutrophil response to *T. gondii* infection of the intestinal epithelium

Phagocytes involved in innate immunity (i.e., dendritic cells, macrophages, and neutrophils) are the first cells to encounter intracellular pathogens after they cross the epithelial barrier of the intestine [61]. In mice, neutrophils are recruited in abundance to the sites of *T. gondii* infection and account for a high proportion of *T-gondii*-invaded-phagocytes in the small intestine following oral entry,[42] **Figure 4Ai**. However, little is known about how neutrophils traffic and behave at the gut-infection site, and data in mouse studies remain to be validated in human models. Therefore, to gain insight into the behavior and response of human neutrophils to *T. gondii* infection we employed our intestinal tissue MPS to assess neutrophil recruitment behavior to the infection site from a neighboring blood vessel, **Figure 4Aii**. We infected Caco-2 intestinal epithelial tubes in our intestinal tissue MPS with ME49 tachyzoites for 48 h, as previously described, **Figure 3B**. Neutrophils isolated from healthy donor blood were introduced to the endothelial lumen of infected systems and observed for migration behavior over a course of 16h. Time-lapse imaging of the intestinal tissue MPS revealed neutrophils migrating within the endothelium and some events of transendothelial migration across the endothelium into the surrounding matrix towards the infected epithelium, **Figure S3 and Video S1-2**. We observed increased neutrophil speed and displacement from the endothelium in our infected systems compared to uninfected controls, **Figure 4B**. Migrating neutrophils were also seen in the matrix near the infected epithelium, and some small fractions of neutrophils were observed within the lumen of the infected epithelium, **Figure 4C**. Confocal imaging also revealed the presence of fluorescently labeled neutrophils within the infected epithelium co-localized with mCherry-tagged ME49 tachyzoites, **Figure 4C (inset)**.

**Figure 4.**
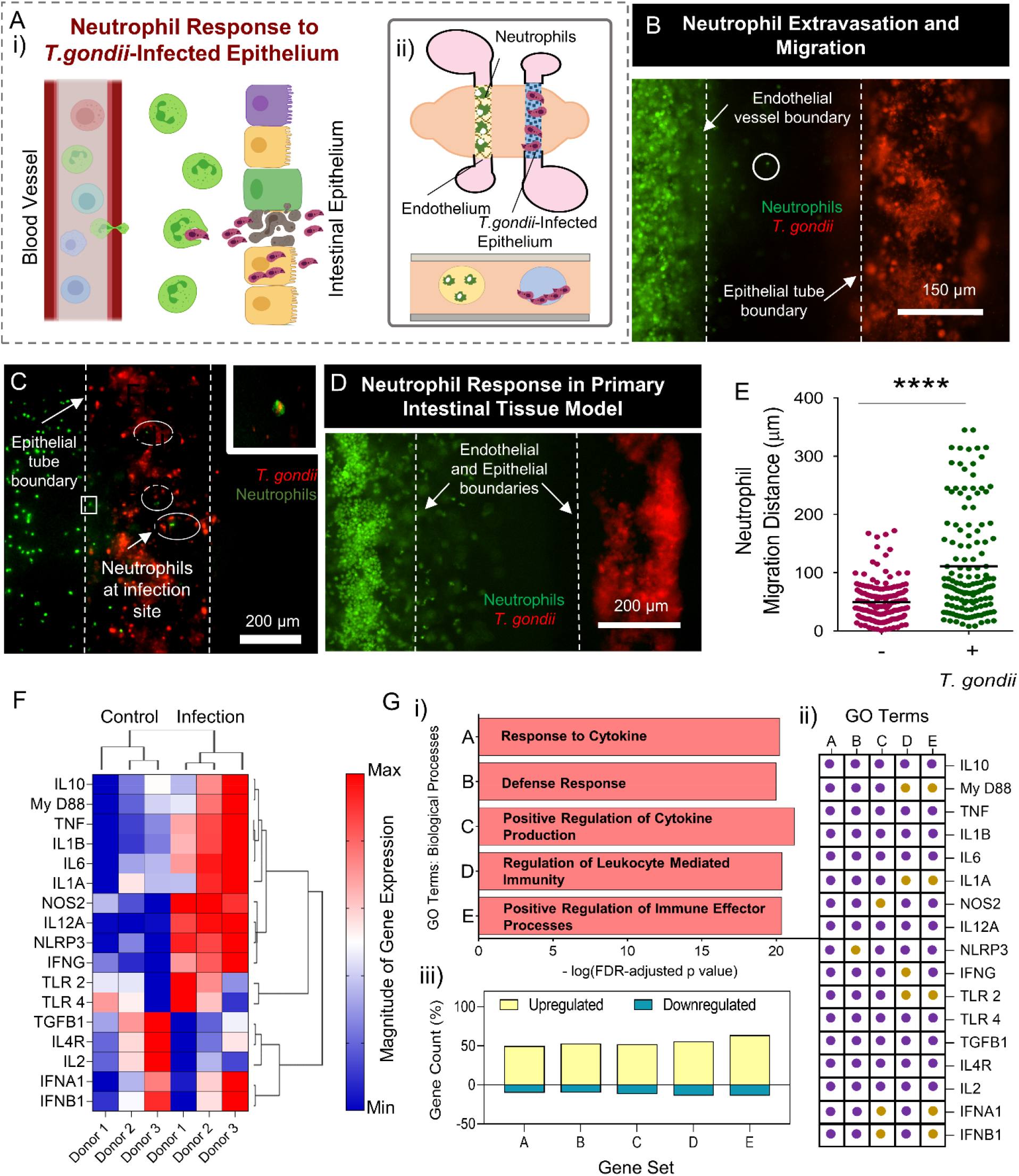
Neutrophil response to *T. gondii* infected epithelium. **(A)**Schematic depicting neutrophil trafficking and interaction with a *T. gondii-*infected intestinal epithelium as seen *in vivo* (i). Illustration representing the spatial distribution of components (a *T*.*gondii-*infected intestinal epithelium, an endothelial vessel and neutrophils) used in the intestinal tissue MPS to explore neutrophil response. (ii). **(B)** Confocal image showing the interface between the gut epithelium and endothelium. Caco-2 epithelial tubes were infected with m-cherry tagged *T. gondii* for 48h before introducing neutrophils into the endothelial vessel. Neutrophils were seen to extravasate and migrate towards the infected epithelium. **(C)** Confocal image showing neutrophils trafficking to the infection site with some neutrophils interacting with the *T*.*gondii*-infected epithelium. White dotted circular lines highlight neutrophil co-localization with the infected epithelium. Inset shows co-localization of neutrophils with *T. gondii*. **(D)** Confocal image showing neutrophil extravasation and trafficking towards a *T. gondii-*infected epithelial tube generated from primary intestinal epithelial cells. Epithelial tubes were infected with m-cherry tagged *T. gondii* for 72 h before introducing neutrophils into the endothelial vessel. **(E)** Grouped scatter plot showing migration distance following extravasation of neutrophils towards *T*.*gondii*-infected primary intestinal epithelium. Each dot represents the migration distance of a single neutrophil from the endothelial vessel boundary (asterisk denotes P value of ≤ □0.0001). **(F)** Differential gene expression and hierarchical clustering analysis of 17 genes conducted in neutrophils from infected systems relative to control systems. Neutrophils from 3 nondiseased donors were used with gene expression profiles analyzed for each donor in control (without infection) and infected systems. Each cell in the matrix corresponds to the expression level of one gene in a sample. The intensity of the color from blue to red indicates the magnitude of differential expression (see color scale to the right of the image). The dendrograms at the top of the figures indicate relationship among experimental conditions which define clusters of conditions (control and infected). The dendrograms at the left of the figures indicate relationship among the profiles of the selected genes which define clusters of higher and lower expression in infected systems, after clustering analysis. Hierarchical clustering was conducted using average linkage clustering with Pearson correlation as the default distance metric. **(G)** Bar graph showing the top five most relevant GO terms associated with the 17 genes analyzed with their corresponding – log(FDR-adjusted P value) (i). GO terms associated with each gene are highlighted with purple dots and genes not-associated are highlighted with light dots (ii). The percentage of genes associated with each GO term according to their fold changes in expression (increase in yellow and decrease in blue) (iii).

When we infected primary small intestinal epithelial cells within the intestinal tissue MPS, a similar increase in end-to-end displacement was observed compared to uninfected systems, **Figure 4D-E**. To directly examine the effects of *T. gondii* infection on gene transcription, we performed RT-qPCR analysis on neutrophils in our model with an infected and uninfected primary small intestinal epithelium. We selected 17 genes that are involved in establishing a protective immunity and activating immune effector functions against invading pathogens. Differential gene expression analysis revealed that infection of the intestinal epithelium with *T. gondii* indeed upregulates multiple genes in neutrophils involved in innate immune responses, including IL10, MYD88, TNF, IL1B, IL6, IL1A, NOS2, IL12A, NLRP3, and IFN-γ, **Figure 4F**. We identified the 10 most significant Gene Ontology (GO) terms related to biological processes, **Table S1**, based on the GSEA molecular signatures database (http://www.gsea-msigdb.org/). The five most relevant GO terms, **Figure 4G**, represent biological processes related to response to cytokines, defense response, positive regulation of cytokine production, regulation of leukocyte mediated immunity and positive regulation of immune effector functions, **Figure 4Gi**. The majority of the genes chosen in our study are involved in these biological processes, **Figure 4G ii** and the overall gene expression of each of these gene sets was generally upregulated in neutrophils from the infected systems, **Figure 4G iii**. These expression patterns suggest that biological processes such as defense response, regulation of leukocyte mediated immunity, and positive regulation of immune effector functions which are key functions of neutrophils during acute *T. gondii* infection are active in neutrophils from our infected systems. Together, these results show that infection of the intestinal epithelium with *T. gondii* in our intestinal tissue MPS elicits responses consistent with those observed *in vivo* behavior where increased transendothelial migration, trafficking of neutrophils towards the infection site, and activation of pathways involved in defense response to invading pathogens are observed.

### IFN-γ mediated neutrophil response to *T. gondii* infection

In the context of *T. gondii* infection, IFN-γ is a cytokine that plays a major role in host resistance and is critical for coordinating protective immunity [62–65]. IFN-γ mediates its protective effects by stimulating lysosomal activity [66], inducing expression of nitric oxide synthase and effector genes [67], and modulating metabolic activity [68] in phagocytes. Neutrophils, upon infiltration to the site of infection, execute a broad range of immune effector functions which include phagocytosis, production of reactive oxygen species (ROS) or antimicrobial peptides, and activation of programmed cell death pathways to reduce the pathogen’s chances of survival [69,70] **Figure 5Ai**. Here, we investigate the influence of stimulation and inhibition of IFN-γ on the effector functions and metabolic activity of neutrophils following an encounter with a *T. gondii* infected epithelium. To evaluate this, we generated tubular epithelium with Caco-2 cells, infected them with *T. gondii* tachyzoites, as previously described, and introduced primary human neutrophils into the lumen of the epithelium, **Figure 5Aii**. As *T. gondii* is an intracellular pathogen, we infected the apical side of the Caco-2 epithelium and allowed incubation for 72 h, with consistent media replenishment, which resulted in some epithelial cells undergoing cell lysis and exposing *T. gondii* to the luminal contents (**Figure 3C)**. Neutrophils, isolated from healthy donor blood, were then added into the epithelial lumen to maximize encounter probability with *T. gondii*, and co-cultured for 6h prior to collection and assessment **Figure 5Bi**. Neutrophil extracellular trap (NET) formation, apoptosis and parasitization of neutrophils were examined, **Figure Bii-iv and Figure S4**. We first examined and quantified the fraction of neutrophils undergoing NET formation, which are released from neutrophils in response to invading pathogens. A key step in NET formation is the release of antimicrobial DNA complexes into the cytosol of the cell. As such, the measurement of decondensed nuclei has been a recognized method for quantifying neutrophils undergoing NET formation [71]. We used Hoechst nuclear stain to distinguish between intact and decondensed nuclei, where intact was characterized by normal trilobed nuclei and decondensed was characterized by diffuse staining with large nuclear area, **Figure 5Ci**. Neutrophils were stained with Hoechst prior to introduction into the lumen of *T*.*gondii* infected epithelium and removed after 6h for analysis. IFN-γ stimulation significantly increased the fraction of neutrophils undergoing NET formation compared to control infection. Conversely, blocking IFN-γ significantly decreased the fraction of NET-forming neutrophils relative to IgG controls, unstimulated controls, and IFN-γ stimulated systems. To evaluate the influence of IFN-γ on apoptotic cell death, we examined the induction of apoptosis using a caspase 3/7 activity assay. Our results indicate that blocking IFN-γ significantly triggered apoptosis while IFN-γ stimulation slightly but not significantly suppressed it, **Figure 5Cii**. We then examined the effects of IFN-γ on the fraction of *T. gondii* parasitized neutrophils. To eliminate parasites non in contact with neutrophils from analysis, neutrophils collected from the epithelium were resuspended in culture medium and centrifuged at 200g for 3 min prior to imaging. Consistent with the trends observed with NET formation, IFN-γ stimulation significantly increased the fraction of *T. gondii* parasitized neutrophils relative to unstimulated systems, **Figure 5Ciii**. Blocking IFN-γ decreased the fraction of *T. gondii* parasitized neutrophils compared to IFN-stimulated systems, demonstrating the sensitivity of neutrophils to IFN-γ stimulation.

**Figure 5.**
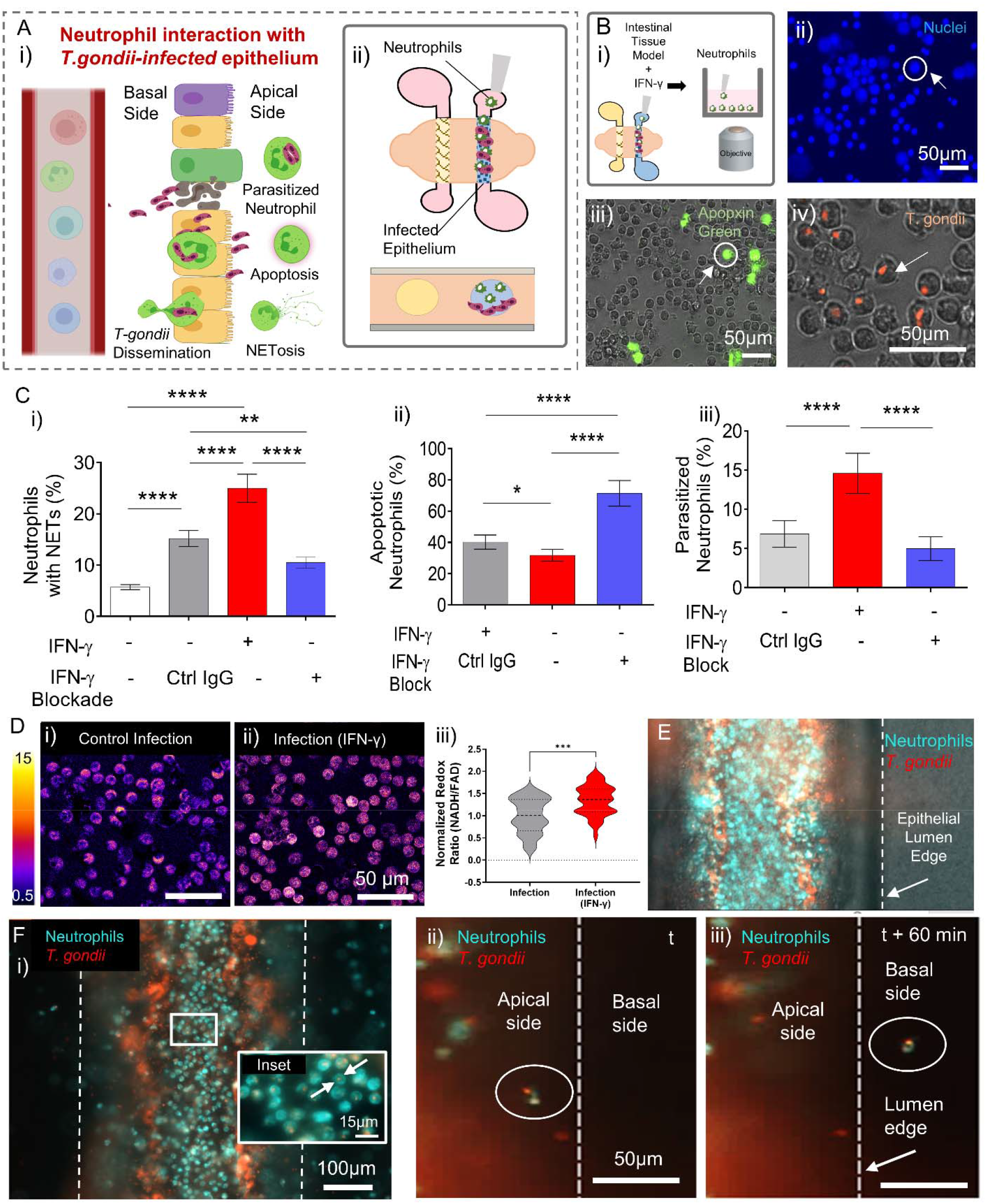
Neutrophil effector functions and contribution to *T. gondii* dissemination beyond the intestinal epithelium. **(A)**Schematic depicting *in vivo* responses of neutrophils at the site of *T. gondii-*infected intestinal epithelium (i). Illustration representing the spatial distribution of components (neutrophils introduced directly into the *T*.*gondii-*infected intestinal epithelium) used in the intestinal tissue MPS to investigate neutrophil effector functions. **(B)** Caco-2 epithelial tubes were infected with m-cherry tagged *T. gondii* for 72h before introducing neutrophils into the lumen of the tube. Neutrophils were co-cultured with the infected epithelium for 6 h prior to collection in well plate and imaging (i). Immunofluorescence image showing neutrophils stained with Hoechst 33342 for DNA. White circular line highlights neutrophils with decondensed nuclei indicating or NET formation (ii). Combined brightfield and fluorescence image showing neutrophils stained with Apopxin Green (abcam) as indicator of cells undergoing apoptosis, white circle highlights an Apopxin Green positive neutrophil (iii). Combined brightfield and fluorescence image showing parasitized neutrophils, white arrow highlights a neutrophil directly interacting with a *T. gondii* parasite (iv). **(C)** Bar graphs showing percentage of neutrophils undergoing NET formation (i), apoptosis (ii) and parasitization (iii) in response IFN-γ stimulation and blockage. Values are presented as mean ±SD of neutrophil response from 3 nondiseased donors (asterisk denotes ****p≤ □0.0001, **p≤ □0.01 and *p≤ □0.05). **(D)** Optical metabolic imaging was used to visualize intracellular NAD(P)H and FAD fluorescence intensities of neutrophils in infected systems without (i) and with IFN-γ stimulation (ii) [redox ratio = NAD(P)H intensity divided by FAD intensity]. Violin plots showing the analysis of neutrophil redox ratio based on NAD(P)H and FAD intensity (iii), (asterisk denotes ***p≤ □0.001). **(E)** Combined brightfield and fluorescence image showing neutrophils within a *T. gondii*-infected epithelium. **(F)** Fluorescence image showing some neutrophils with internalized *T. gondii* after 6h co-culture with infected epithelium(i). Fluorescence images showing *T. gondii* trafficking by neutrophils across the epithelial barrier, white dashed line indicates the epithelial boundary separating apical and basal surface (ii-ii).

We also investigated the influence of IFN-γ on neutrophil metabolic activity by means of optical metabolic imaging, which quantifies relative amounts of reduced NADH and FAD [72,73], and by extension the redox ratio. The optical redox ratio is used to obtain information on the dynamic changes in oxidation-reduction rates in cells and is sensitive to alterations in cellular metabolic rates [72]. Optical NADH/FAD redox ratios increased in neutrophils after 6h in our IFN-γ stimulated systems indicating increased metabolic activity, **Figure 5D**, and these observations were conserved when the analysis was limited to *T. gondii* internalized neutrophils, **Figure S5**. Relative to unstimulated systems, lower FAD intensity was observed in response to IFN-γ stimulation. Altogether, these findings suggest that neutrophil effector functions are IFN-γ-dependent where IFN-γ stimulation enhances neutrophil effector functions by increasing events of *T*.*gondii* internalization, NET formation, and cellular metabolic activity of neutrophils while prolonging their life span. More importantly, these results demonstrate the utility of our intestinal tissue MPS in testing the sensitivity of immune cells to stimulants that may influence their response to pathogens.

Following an encounter at the infection site, engulfed microbes are carried by migratory phagocytes such as macrophages and neutrophils beyond the epithelia of barrier organs into deeper tissue and draining lymph nodes, contributing to pathogen dissemination to other organs, [74] **Figure 5A**. In *T. gondii* infected mice, parasite-containing neutrophils in the small intestine have been shown to transport the parasite across epithelial barriers to facilitate parasite spread both within the intestine to other regions and beyond to the spleen and mesenteric lymph nodes [42,75,76]. To investigate whether dissemination of *T. gondii* by migratory neutrophils beyond the epithelium could be observed in our intestinal tissue MPS, we introduced neutrophils into an infected intestinal epithelium generated from Caco-2 cells **Figure 5E**. Within 6 hours, *T. gondii* containing neutrophils could be seen within the lumen of the intestinal epithelium, **Figure 5Fi**. Time course imaging revealed translocation of *T. gondii* containing neutrophils from the apical side to the basal side of the epithelium, **Figure 5F ii-iii**. These results show that our intestinal tissue MPS can recapitulate the mechanisms of parasite dissemination by migratory phagocytes as seen *in vivo*. Furthermore, our ability to visualize immune cell-mediated transport of pathogens within our intestinal tissue MPS could provide valuable insight into how intracellular pathogens disseminate and present opportunities to investigate new targets for therapeutic intervention.

### Innate Immune cell response to parasite invasion and replication in the intestinal epithelium

The innate immune system coordinates the first immunological defense against an invading pathogen [77]. As cytokine production by intestinal epithelial and endothelial cells is a key feature of early host immune response, we wanted to evaluate the capacity of our intestinal tissue MPS to produce the inflammatory mediators required for immune effector functions in response to parasite invasion, **Figure 6A**. We analyzed inflammatory cytokines secreted in culture media prior to the addition of immune cells using a Luminex multiplex bead-based ELISA assay. Our results revealed that infection of the primary small intestinal epithelium with *T. gondii* induced significantly higher levels of proinflammatory cytokines/chemokines MCP-1, MIP-1α, IL-1α, IL-1β, GM-CSF, IL-6, and IL-8 secretion in culture media at 48 hpi compared to uninfected controls, **Figure 6B**. Secretion of anti-inflammatory cytokines such as IL-4, IL-10, and IL-13 on the other hand showed no change in the infected systems compared to control uninfected systems. Together, these cytokines make up critical components of the inflammatory milieu that contribute to coordinated immune defenses like immune cell trafficking, activation, and effector functions against parasites during acute infection.

**Figure 6.**
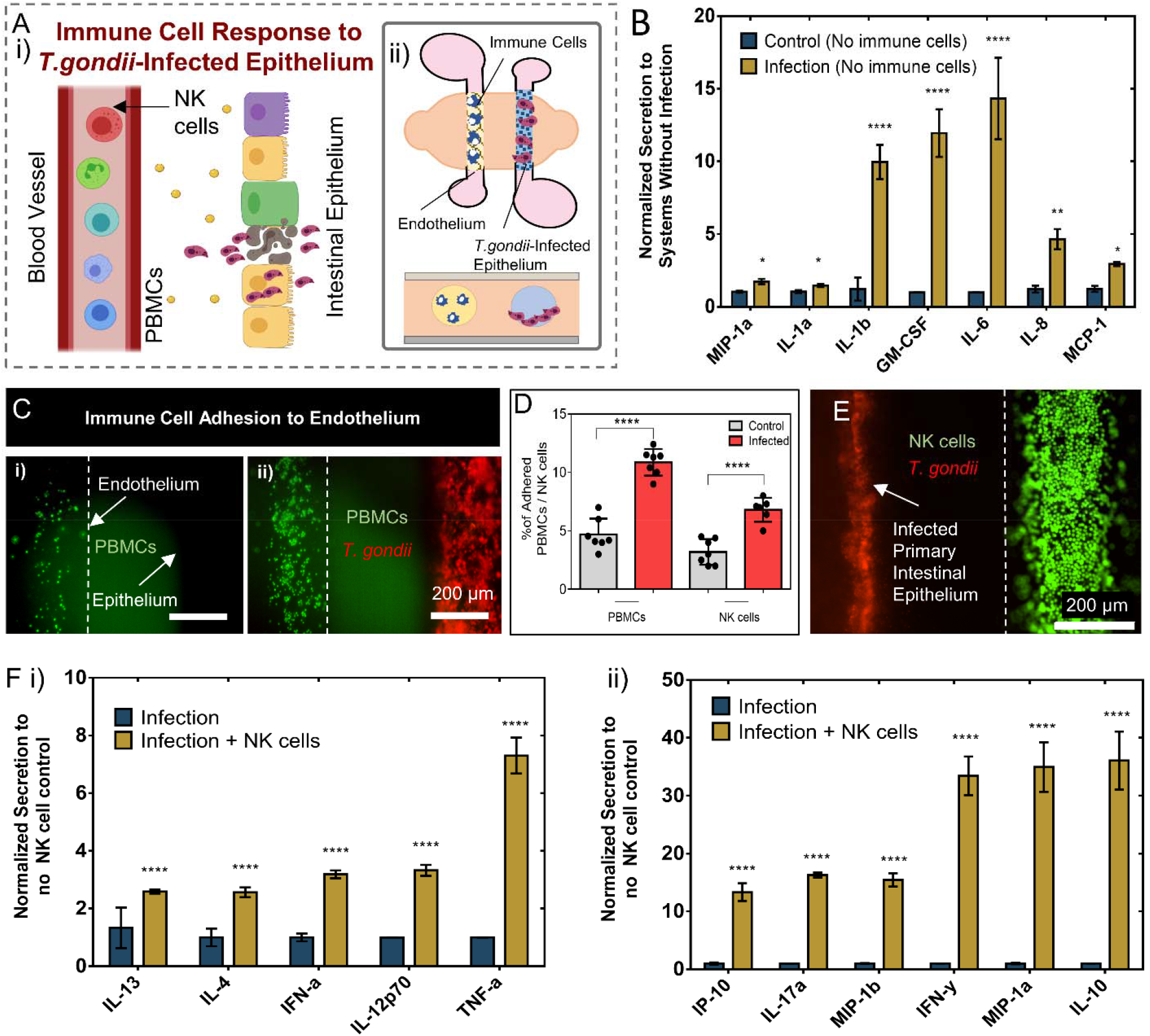
Immune cell response and cytokine secretions within the intestinal tissue MPS. **(A)**Schematic conceptualization of immune cell response to soluble factor signaling during the initial stages of *T. gondii-*infection (i). Illustration representing the spatial distribution of components (*T*.*gondii-*infected intestinal epithelium, endothelial vessel, PBMCs or NK cells) used in the intestinal tissue MPS to investigate immune response. **(B)**Cytokine concentrations measured in media collected from the intestinal tissue MPS consisting of endothelial vessel and primary intestinal epithelial tubes infected with *T. gondii* normalized to control, uninfected systems (asterisk denotes ****p≤ □0.0001, **p≤ □0.01 and *p≤ □0.05). **(C)** Fluorescence image showing differences in PBMC adhesion events to the endothelial vessel in control (i) and infected (ii), Caco-2 epithelium. **(D)** Bar graphs showing percentage of PBMCs and NK cells adhering to the endothelium of control and infected systems following co-culture for 2h. PBMCs and NK cells were added to the lumen of the endothelial vessel after Caco-2 epithelial infection by *T. gondii* for 48h. Five devices were prepared on two different days for each condition. **(E)** Fluorescence image showing *T. gondii*-infected primary intestinal epithelial tube in co-culture with an endothelial vessel containing NK cells. Epithelial tubes were infected for 72h prior to adding NK cells into the endothelial vessel. **(F)** Cytokine concentrations measured in media collected from infected intestinal tissue MPS consisting of an endothelial vessel, NK cells and primary intestinal epithelial tubes infected with *T. gondii* normalized to infected systems without NK cells. In all cytokine measurement experiments nine devices were prepared on two different days for each paired conditions (infected vs. control, and infection in the absence vs. presence of NK cells), media from three devices were pooled to make one replicate. Values are presented as mean ±SD asterisk denotes ****p≤ □ 0.0001).

Adhesion molecules on endothelial cells are known to mediate leukocyte rolling, adhesion, and subsequent transendothelial migration as part of an inflammatory response to injury or infection. We investigated the effects of infection on the firm adhesion of immune cells to the endothelium. Fluorescently labeled primary human peripheral blood mononuclear cells (PBMCs) were added to the endothelium 48 hpi with ME49 *T. gondii* and washed away after 2 hours to evaluate endothelial adhesion of immune cells, **Figure 6C**. Indeed, the percentage of PBMC adhering to the endothelium doubled relative to systems without infection, **Figure 6D**. A similar increase in endothelial adhesion was also observed when NK-92 cells were added to the endothelial vessel, **Figure 6D and Figure S6A**. Overall, these results provide strong evidence of epithelial-endothelial crosstalk during parasite infection that promotes immune cell adhesion to endothelium, a critical step in endothelial transmigration and recruitment of immune cells to the infection site. Concurrent with the high levels of proinflammatory cytokines observed in our system, increased levels of soluble adhesion molecules, sP-selectin, sICAM-1, and sE-selectin were also found in the culture media of infected systems, **Figure S6B**.

Before adaptive immunity is established, NK cells restrain the spread of infection by secreting inflammatory cytokines that are critical for stimulating protective immunity[78]. To explore NK cell-mediated production of cytokines in our *T. gondii* infected intestinal tissue MPS, we compared cytokine secretions in infected systems in the presence and absence of NK cells. NK-92 cells were introduced in the endothelial lumen of infected systems at 48 hpi and cultured for 24 hours, **Figure 6E**. Secreted factor analysis was performed on media collected from systems 24 hours after the introduction of NK cells into the lumen of the endothelium. Our results show that the presence of NK cells in infected systems led to significantly higher levels of pro-inflammatory factors including IFN-α, IL12p70, TNF-α, IP-10, IL-17a, MIP-1α, MIP-1β, and IFN-γ, **Figure 6F**. The presence of NK cells in infected systems also induced significantly higher secretion of IL-10, IL-4, and IL-13 which are known to suppress pro-inflammatory cytokine production. While the induction of a robust inflammatory response is critical for host resistance against *T. gondii*, unregulated inflammation can result in exacerbated immunopathological reactions causing tissue damage. Thus, anti-inflammatory factors may have critical roles in reducing inflammatory reactions and suppressing collateral damage because of upregulated inflammation during parasite infection.

## Discussion

Development of improved treatment strategies against parasitic infections requires increased knowledge of the pathogen-tissue-immune system interactions. Murine models have made substantial contributions to our understanding of innate immunity against parasitic infections, but sufficient differences in the organization of the immune system between humans and mice warrant the need for improved models. Therefore, the development of new tools that allow researchers to elucidate mechanisms involved in immune responses and defense strategies against parasites and other activators of the immune system in humans could accelerate the development of antiparasitic drugs and therapeutic strategies. In this regard, we have bioengineered an integrated microphysiological system of the intestinal tissue for exploring human innate immune cell responses to parasite infection in the gut. Tubular tissue structures like the intestines and the vasculature system serve essential roles in separation of internal contents and pathogenic microbes in the intestinal lumen from systemic dissemination. Our human intestinal tissue MPS incorporates *in vivo*-like tubular geometries of the intestinal epithelium and endothelium allowing us to explore host-parasite interactions and tissue-dissemination that occur during the initial stages of infection. Using our model, we examined epithelial infection by *T. gondii*, an intracellular parasite that initially infects the small intestine following ingestion. We introduced a suspension of *T. gondii* tachyzoites directly onto the apical surface of the intestinal tubes and using live-cell microscopy we demonstrated that our tubular intestinal epithelium supports invasion and intracellular replication of *T. gondii*. As an important component of the intestinal immune network, the innate immune system is considered the most important defense line against pathogens breaching the epithelial barrier and plays a pivotal role in maintaining immunity and preventing tissue dissemination through the vascular system. Although incorporating tubular intestinal epithelium within scaffolding matrices has previously been reported [25,29,46], the originality of our model relies on the integration of vascular and immune cell components. This enables us to investigate the various facets of innate immune cell responses, including extravasation and trafficking, to *T. gondii* infection of the intestinal epithelium and its translocation across the epithelial barrier. One additional design aspect of our system is the compatibility with live-cell microscopy enabling dynamic visualization of host-microbe interactions occurring at the interface between the intestinal epithelium and the neighboring vasculature at high spatiotemporal resolution. Using these techniques and molecular analysis, we examined early responses from human innate immune cells like neutrophils and NK cells and investigated parasite-immune cell interactions during epithelial infection that influence immune effector functions and tissue dissemination.

Current *in vitro* studies of parasitic infections primarily rely on human cell lines that often do not recapitulate *in vivo* phenotypes. On the other hand, maintaining long-term proliferative cultures of the human intestinal epithelium *in vitro* have met with continuing difficulty due to the complex interactions between cell types and the presence of molecular signaling required for stem cell maintenance. Organoid technologies enable the expansion of primary intestinal stem cells and are regarded as a powerful *in vitro* tool for modeling intestinal epithelial tissue due to their structural and functional resemblance to *in vivo* tissue [79–81]. While apical access can sometimes be limited when using these techniques, methods to form epithelial monolayers from intestinal organoid fragments have been reported [82–84], where the major epithelial cell types found in 3D organoids are retained when cultured on hard surfaces coated with Matrigel or collagen [85]. Building on these methods, here we adapted our intestinal tissue MPS to integrate the formation of tubular epithelium using primary human intestinal epithelial cells.

We used a simple micromolding based technique to generate tubular structures within a hybrid matrix, composed of type I collagen, type IV collagen, and Matrigel. Our data indicates this matrix-supported culture and differentiation of human intestinal epithelial cells, both cell lines (e.g. Caco-2) and primary organoid-derived intestinal stem cells obtained from jejunal tissue resections. Maintenance of primary human-patient derived small intestinal cells also require stimulation from the basal compartment by niche factors that help retain stem cell components while supporting differentiated enterocyte-goblet cell function [50,80]. To support the integration of the endothelial vessel and immune cells (i.e., neutrophils, NK cells, and PBMCs) into our systems, we optimized culture protocols and media compositions for differentiating a primary intestinal epithelium and to support all cell types within the intestinal tissue MPS. Although several molecular and functional properties of the intestinal epithelium within the system are expected to change due to constantly evolving differentiation states and influence from co-culture with other cell types, we confirmed through visual inspection of the system the epithelium and endothelium remain intact. Further, we show that phenotypic characteristics and barrier properties of both the epithelium and endothelium are retained for the duration of the experiments (5 days post endothelial cell seeding), which correlates with the turnover of intestinal epithelia *in vivo [86]*. Intestinal tubes generated within the hybrid matrix, both from the Caco-2 cell line and primary intestinal stem cells, exhibited key properties of the intestinal epithelium including a polarized epithelium expressing markers of tight junctions, goblet cells, and microvilli formation. Interestingly, gene expression analysis showed that expansion of primary intestinal stem cells in tubular structures resulted in increased maturation and differentiation, while retaining key characteristic features of the jejunal epithelium compared to primary epithelial monolayer formed in transwells. Gene expression analysis also demonstrated upregulation of pathways involved in gut epithelium regeneration compared to transwell cultures. These results agree with previous reports showing that a suitable matrix with relevant ECM proteins and biophysical properties are key factors in providing physical support and facilitating the differentiation of intestinal stem cells [58].

By integrating blood vessels, circulating innate immune cells, and primary intestinal epithelial cells within a biomimetic 3D scaffolding matrix, we incorporated key cellular components and relevant cell-ECM interactions of the intestinal tissue for improved modeling of parasite interaction with the diverse cell types of the epithelium and immune cell responses during epithelial infection. However, one limitation in this work is that we did not incorporate components of the microbiome that can also impact the mechanisms of parasite interaction with the epithelium and innate immune cells. This would require additional control over environmental factors such as oxygen levels to accommodate oxygen-sensitive species of the microbiome. Future studies could incorporate this layer of complexity by incubating the intestinal tissue MPS in a hypoxia incubator (i.e., 1% O2) and perfusing oxygenated medium through the vascular lumen.

Initial responses against *T. gondii* are managed by innate immune cells, with neutrophils, dendritic cells, and monocytes playing a central role [8,87,88]. Early depletion studies in mice using monoclonal antibodies showed that neutrophils are important for host survival during acute infection [89]. Further highlighting their importance, depletion of neutrophils showed decreased levels of gamma interferon (IFN-γ), interleukin-12 (IL-12), and tumor necrosis factor (TNF), indicative of a weaker type I immune response, and development of lesions in multiple organs including lung, liver, and brain. Using our intestinal tissue MPS, we recapitulated events with human neutrophils including extravasation and migration towards the *T. gondii* infected epithelium, reminiscent of the process of leukocyte trafficking to the site of infection previously observed in mouse models [42]. To reduce donor-to-donor variability from adult intestinal stem cells, we used organoid-derived stem cells from a single donor for generating tubular intestinal epithelium, however, neutrophils from multiple blood donors were used to capture responses from neutrophils in this study. Our results showed that the interaction of *T. gondii* infected primary small intestinal epithelium with neutrophils led to increased expression of genes involved in biological processes including response to cytokines, defense response, positive regulation of cytokine production, regulation of leukocyte mediated immunity, and positive regulation of immune effector functions. However, in this work, we used RT-qPCR to analyze ≈17 genes related to neutrophil response, which only represents a fraction of the total number of alterations involved in establishing this immune response. Future investigations could include RNA-sequencing to provide a more in-depth molecular analysis of the biological processes activated in neutrophils in response to *T. gondii* infection in our model. On the other hand, our study also highlighted changes in the functional behavior of neutrophils that correlate with the biological process involved as determined by gene expression analysis. In line with the transcriptional changes observed, dynamic visualization of neutrophil behavior at the epithelial and vascular interface showed increased migration speed and end-to-end displacement from the endothelium in response to the infection of the epithelium. Strikingly, some instances of direct interaction between neutrophils and excysted *T. gondii* within and near the epithelium could also be observed, demonstrating the capability of our model in capturing parasite invasion, replication, and initial response of neutrophils in the same experiment.

Host-directed therapies that target signaling pathways for parasite clearance bypass many problems encountered by anti-parasitic drugs including poor patient compliance and the emergence of drug-resistance parasites [90]. By targeting host pathways that are redundant in the host but are critical for parasite survival, there is a reduced chance of developing treatment resistance due to the slow rates of mutations in molecules and processes in the host relative to parasites. In this work, we examined the effects of a host-targeted pathway on the effector functions of neutrophils. Once at the site of infection, local production of inflammatory mediators regulates intercellular communication that mobilizes several defense mechanisms in tissue-resident cells and immune cells [91]. IFN-γ has a multitude of immunomodulatory functions and is considered one of the most potent pleiotropic cytokines [64,65,92]. IFN-γ interactions with T cells, NK cells and activated macrophages have been widely researched, however, investigations into the role of IFN-γ during the initial responses of neutrophils and other innate immune cells to infection by parasites have been limited. We demonstrated the influence of IFN-γ on traditional neutrophil functions including parasitization, NET formation, and apoptosis of neutrophils. IFN-γ treatment favorably affected neutrophil functions with increased parasitization, NET formation, and overall metabolic activity in neutrophils. In contrast, neutrophil apoptosis, the process of programmed cell death, decreased which limits exposure of destructive neutrophil productions to surrounding tissue. IFN-γ blockage reversed these trends, further highlighting the importance of IFN-γ on traditional neutrophil functions of anti-microbial activities and showing the utility of our model for investigation of human immune responses. We also demonstrate the ability of our model to investigate parasite dissemination facilitated by neutrophils, both within the lumen of the intestinal epithelium and beyond by crossing the epithelial wall into the surrounding tissue. These observations are also consistent with data in mice that show the involvement of neutrophils in the spread of *T. gondii* infection with the intestinal tissue [42]. Altogether, we demonstrate the sensitivity of our system in capturing the changes in various effector functions and antimicrobial activity of neutrophils in response to inflammatory stimulus and blockage.

Our secreted factor analysis revealed that our intestinal tissue MPS produces a wide array of proinflammatory cytokines/chemokines (MCP-1, MIP-1α, IL-1α, IL-1β, GM-CSF, IL-6, and IL-8) that enhance and activate defense mechanisms in innate immune cells during *T. gondii* infection. In mice, these factors are known to have immunoregulatory roles during acute infection. For instance, the production of MCP-1 and MIP-1α in our system is consistent with *in vivo* data in mice showing higher expression of these chemokines in *T. gondii* infected intestinal epithelial cells [93]. Also, mice deficient in MCP-1 and its chemokine receptor CCR2 fail to generate adequate immune response during acute *T. gondii* infection [94], highlighting its critical role in regulating innate immunity. Elevated levels of IL-1α and IL-1b have been previously observed in the serum of *T. gondii* infected mice (one week post-infection) and are known inflammatory mediators/regulators of host tissue homeostasis during acute infection [95,96]. Moreover, aside from their involvement in inflammation, chemokines like GM-CSF, IL-6, and IL-8 play important roles in the recruitment and enhanced survival of phagocytic cells like neutrophils during acute innate response to *T. gondii* infection [97–99]. Together, these cytokines make up critical components of the inflammatory milieu that contribute to coordinated immune defenses like immune cell trafficking, activation, and effector functions against parasites during acute infection. In accordance with the increased levels of these cytokines in our infected systems, we observed increased neutrophil trafficking and transcriptional changes associated with responses to cytokines and positive regulation of immune effector functions in neutrophils. While we have not explored the direct influence of these cytokines on the functional behavior of neutrophils, future studies using our model could explore spatial and temporal relationships between neutrophils and infection-derived factors that enhance their overall effector functions against parasites.

During innate immune responses, *T. gondii* infection-induced production of inflammatory cytokines, drive NK cells to produce a host of factors such as IFN-γ and TNF-α that amplify effector functions and inflammatory responses driven by other innate immune cells [100]. Intriguingly, NK cell production of immunosuppressive cytokines that counterbalance inflammatory responses during disseminated pathogenic infection with *T. gondii* have also been found [101], shedding light on the role of NK cells in facilitating immunoregulation. While the mechanisms haven’t been clearly defined, the immunoregulatory roles of NK cells are likely important for preventing inflammation-dependent pathology during parasite infection. Further exploration of secreted factor analysis shows that incorporating NK cells into our systems upregulates several immunoregulatory signaling factors critical for amplifying the production of inflammatory and enhancing effector functions in other innate immune cells. In the presence of NK cells, several cytokines were uniquely elevated IFN-α, TNF-α, IFN-γ, IL-17a, IP-10, IL-4, IL-13, IL-12p70, and IL-10 in our infected systems. IFN-α, a member of the type I interferon family, has recently been shown to enhanced cytokine secretion and cytotoxic potential in NK cells [102], and TNF-α together with IFN-y are major proinflammatory cytokines produced by NK cells as means of host protection against *T. gondii [44]*. Moreover, these two cytokines are recognized as essential immune effectors against intracellular pathogens. IL-17 has been implicated in resistance to *T. gondii* where IL-17R-/-mice showed defects in neutrophil recruitment and increased parasite burden [103]. As NK cells are often considered to function at the interface of the innate and adaptive immune response [104], NK cell production of chemokine, IP-10, can be critical for generating an adaptive immune response and influencing the recruitment of effector T cells to the *T. gondii* infection site [105]. Interestingly, cytokines typically involved in the resolution of cell-mediated inflammation (IL-4, IL-13, and IL-10 [106]) were also elevated in our infected systems with NK cells. Conventionally, IL-4 and IL-13 are thought to counter NK cell effector functions by limiting the production of IL-12. However, recent work shows that stimulation with IL-4 or IL-13 enhances TNF-α and IFN-γ production in NK cells in the presence of IL-12, underscoring the immunoregulatory roles of these cytokines [107]. Elevated IL-10 levels observed are also in line with previous reports showing that NK cells produce IL-10 during acute *T. gondii* infection which interferes with the activation of the adaptive immune responses [108], further highlighting the immunoregulatory roles of NK cells. While the source of these cytokines is unclear in our analysis, secretion of these factors is markedly higher in our infected intestinal tissue MPS with NK cells, indicating a direct or indirect contribution from NK cells. The local effects of this inflammatory environment on the effector functions of neutrophils and other innate immune cells may be tested with future experiments within our platform and using tools/methods for examining complex interactions *in vitro*. Collectively, our secreted factor analysis demonstrated a wide array of immunoregulatory cytokines, chemokines, and adhesion molecules generated within our primary intestinal tissue MPS during *T. gondii* infection, which are essential for generating a robust immune response. These molecules are critically involved in innate immunity against intracellular pathogens in the following ways: 1) promoting leukocyte adhesion to the endothelium and transendothelial migration, 2) trafficking, recruitment, and survival of phagocytic cells, and 3) enhanced activation, cytotoxic activity, and effector functions of immune cells involved in the innate and adaptive immune responses. These data confirm the presence of an immunomodulatory cytokine milieu that can support mounting a strong immune response to parasitic infections.

The primary objective of this work was to create a microphysiological system that recapitulates complex interactions between key components of the human intestinal tissue and to demonstrate its usefulness in studying innate immune responses to parasitic infection. Recreating the initial stages of parasitic infection by *T. gondii* and the accompanying innate immune responses of the intestinal tissue highlights the potential of our microphysiological model to fill the gaps in studying human tissue responses through mechanistic correlations between various molecular and functional cell data to both animal and human clinical data. The readily accessible approach to generating 3D geometries of tubular organs within biomimetic ECM scaffolds, which accommodate the development and maturation of stem cells, and enables dynamic visualization of host-microbe interactions at a high spatiotemporal resolution, could answer questions that have so far been difficult to address and may have substantial potential for discovery of cell-mediated immune responses and development of host-directed therapies.

## Materials and Methods

### Intestinal tissue MPS fabrication and assembly

The intestinal tissue MPS was fabricated using the LumeNEXT approach as previously reported[109]. Briefly, this approach uses a micromolding technique to fabricate one or more lumen structures with variable size, configuration and lumen spacing controlled by micromold design in soft lithography. Here, we use a two-lumen setup, each having a separate inlet and outlet port within a single chamber was used accompanied by perpendicularly oriented side ports. Two stacked PDMS layers, with microscale features patterned into them, formed the culture chambers in which scaffolding ECM gel can be loaded, while removable PDMS rods formed the hollow lumen structures surrounded by the ECM gel. The master molds for the PDMS layers were made using SU-8 100 (Microchem) which were spin-coated onto wafers, soft-baked (i.e., heat at 65°C for 30-40 min and then at 95°C for 90-120 min depending on layer thickness), exposed to UV through a mask of desired patterns and post-baked at 95 °C for 20-30 min. This procedure was repeated for additional layers prior to development in propylene glycol monomethyl ether acetate (Millipore Sigma). After developing, PDMS (Sylgard 184 Silicon Elastomer Kit, Dow Corning Corporation) was applied to the masters at a ratio of 10:1 base to curing agent and allowed to polymerize for 4 h at 80 °C. The rods were drawn from needles with gauge size, 23 gauge (340 µm inner diameter). Prior to device assembly, the PDMS layers and rods were soaked in 100% ethanol for several days to extract any uncured PDMS oligomers. Following PDMS extraction, the rods were placed in between two layers, across the body of the chamber (3 mm in length) in ledge features stemming from the smaller inlet and larger outlet ports to hold the rods in the middle of the chamber. The side ports (4 mm apart) of the chamber were used to fill the chamber with ECM gel and the height of the chamber was about (1.25 mm). Once assembled, the PDMS layers were oxygen-plasma bonded onto a glass-bottom MatTek dish using a Diener Electronic Femto Plasma Surface System.

### ECM gel preparation and loading

The bonded devices were UV sterilized for 20 min and moved to the biosafety cabinet, prior to ECM gel loading. To promote matrix adhesion to PDMS, the device chambers were treated with 1% polyethylenimine (Millipore Sigma) in DI water solution for 10 min, followed by a 30 min treatment of 0.1% glutaraldehyde (Millipore Sigma) in DI water solution. Following surface treatment, devices were flushed with DI water solution 5 times to remove excess glutaraldehyde. A high concentration rat tail collagen I (Col-I) (Corning) neutralized with 0.5N sodium hydroxide (Fisher Scientific), was mixed with 7.5 pH 5X phosphate buffered saline (PBS), complete growth medium or organoid expansion medium, human placental type IV collagen (Col-IV) and Matrigel to achieve a final ECM solution containing 4 mg/mL Col-I, 15% Matrigel and 50μg/mL Col-IV. The pH of the ECM mix was adjusted to 7.2 pH prior to loading the mix into the gel-chamber of the device. The devices were first kept at room temperature for 20 min then moved to an incubator at 37°C for at least 1 h prior to cell loading. To prevent dehydration during polymerization, PBS was added to the MatTek dish surrounding the devices. PDMS rods were then removed leaving behind hollow tubular structures within the ECM gel which can be lined with cells.

### Crypt Isolation and gut organoid culture

Small intestinal crypts were isolated using a previously established protocol.[110] from jejunal tissue resection samples removed from nondiseased tissue of de-identified individuals[110]. Jejunal tissue resections were performed at University of Wisconsin-Madison upon the donors’ informed consent and methods were carried out in accordance with Institutional Review Board (IRB) (Protocol# 2016-0934). Small intestinal organoids were established from isolated crypts by resuspending in Matrigel (Corning; growth-factor-reduced, phenol-red-free formulation), and culturing in 24-well plates (Polystyrene, Nunc, Non-Treated Multidishes, Thermo Fisher Scientific) at 37°C in the organoid expansion media. Expansion medium consisted of a mixture of base medium (BM;45%v/v), L-WRN conditioned medium (CM;45%v/v), supplement mix, 1× Primocin (InvivoGen) and 10% heat-inactivated fetal bovine serum (FBS; Millipore Sigma). BM was prepared from Advanced Dulbecco’s modified Eagle’s medium (DMEM)/F12 (Thermo Fisher Scientific) supplemented with Glutamax (2 mM; Thermo Fisher Scientific), Hepes (10 mM; Thermo Fisher Scientific), penicillin/streptomycin (Pen/Strep) (Thermo Fisher Scientific), B-27 Supplement (Thermo Fisher Scientific), N2 supplement (Thermo Fisher Scientific) and 1% cell-culture grade bovine serum albumin (BSA). L-WRN CM was prepared using L-WRN cells from American Type Culture Collection (ATCC, catalog no. CRL-3276) and a previously published protocol [111]. The supplement mix consisted of EGF (50 ng ml−1; Peprotech), N2 supplement (Thermo Fisher Scientific), human [Leu15]-gastrin I (10 nM; Millipore Sigma), N-acetyl cysteine (500 μM; Millipore Sigma), nicotinamide (10 mM;Thermo Fisher Scientific), A83-01 (500 nM; Tocris), SB202190 (10 μM; Selleckchem), prostaglandin E2 (10 nM; Tocris), Y-27632 (10 μM; Seleckchem) and CHIR99021 (5 μM;. Tocris). Jejunal organoids were split every 7 to 9 days as previously described.[112] Based on this established protocol, Matrigel plugs containing organoids were digested in 0.5 μM ethylenediaminetetraacetic acid (EDTA; Invitrogen) and collected. After centrifugation at room temperature for 3 min at 300×g, supernatant was removed followed by incubation in trypsin (Sigma Aldrich) for 2 min at 37 °C. Trypsin was neutralized using Advanced DMEM/F12 supplemented with 20% FBS and 1% Pen/Strep followed by mechanical dissociation of organoids by vigorous pipetting (about 30 times) to make small organoid fragments. After centrifugation at room temperature for 3 min at 300×g, pellets were reconstituted in 50:50 mix of fresh Matrigel and expansion medium at 1:3 to 1:4 ratio and cast in 25-μL droplets in 24-well plate. Matrigel droplets containing organoid fragments were then polymerized for 15 min at 37 °C before adding 500μL expansion media to each well. Expansion media was changed every 2 to 3 days and organoids were used between passage 5 - 25. Y-27632 was only used for the first 48 h after single-cell dissociation to prevent detachment-induced cell apoptosis.

### Caco-2 epithelial tube formation

Caco-2 cell line was acquired from the ATCC and maintained in Eagle’s Minimum Essential Medium (EMEM, Millipore Sigma) supplemented with 20% FBS and 1% Pen/Strep. To generate tubular epithelium, Caco-2 cells were detached using a trypsin/EDTA solution and resuspended at 17 million cells per mL of supplemented EMEM. After removal of the PDMS rod 3μL of cell suspension was introduced into the hollow tubes through the inlet ports and cells were allowed to fill the lumen structures by passive-pumping. The microdevice was rotated every 30 min over a 2-hour period using a previously described method [109] to line the hollow tube with epithelial cells. Non-adherent cells were washed off with culture medium and excess medium was added to the inlet/outlet ports. After completion of tube formation (∼24-48h) the epithelial tubes were cultured in supplemented EMEM medium with reduced FBS to 10% for 5 - 7 days prior to co-culture with endothelial vessels.

### Primary intestinal epithelial tube formation

Jejunal organoids were collected at day 7 to 9 after passaging and dissociated into small organoid fragments as described above. Pellets of organoid fragments were resuspended in expansion medium at a density of 5 million cells per mL. As previously described 3μL of organoid cell suspension was introduced into the hollow tubes through the inlet ports. Cells were allowed to settle on the bottom-half of the tubular structure for 24h at 37 °C in 5% CO_2_ after expansion medium was added to the gel ports and the inlet/outlet ports during culture. After 24h non-adherent cells were gently washed with expansion medium. Fresh organoids fragments were prepared and seeded again into the tube. Expansion medium was added to the ports and the microdevice cultured for an additional 24h while flipped upside-down to facilitate adhesion of cells to the top-half of the hollow tubular structure. After completion of tube formation (∼72h) the primary intestinal epithelium was cultured in differentiation medium (DM) for 5 - 7 days. For differentiation, the culture medium was replaced with BM (80%v/v), L-WRN CM (10%v/v), supplement mix (without Y-27632 and reduced CHIR 99021 to 500nM), 1× Primocin (InvivoGen) and 10% heat-inactivated fetal bovine serum (FBS; Millipore Sigma). DM was changed twice daily. After 5 - 7 days of differentiation, human umbilical vein endothelial cells (HUVECs) (Lonza) were added to the adjacent hollow tube to generate co-cultures of epithelial and endothelial tubes.

### Co-culture of epithelial tubes with endothelial tubes

HUVECs were maintained in EGM-2 MV Bulletkit medium (Lonza) and used until passage 8. To generate co-culture of primary intestinal tubes with endothelial vessels, HUVECs were detached using a trypsin/EDTA solution and resuspended at 15 million cells per mL of modified DM. Modified DM was made by replacing BM in the DM formulation with EGM-2 MV Bulletkit medium. 3μL of endothelial cell suspension was introduced into the cell-free hollow tube next to the epithelial tube and device was rotated every 15min over a 2-hour period as previously described. Modified DM was added to the gel ports and epithelial inlet/outlet ports during this process to nourish the epithelial cells. After lining the tube with endothelial cells, nonadherent cells were gently washed with modified DM. The epithelial and endothelial tube co-cultures were maintained in modified DM for the remainder of the experiments unless otherwise stated. To support co-culture of HUVEC vessels with Caco-2 epithelial tubes, the supplemented EMEM medium with reduced FBS was mixed with EGM-2 MV at 50:50% v/v dilution and used as culture medium for the remainder of the experiments.

### Co-culture of epithelial and endothelial tubes with NK cells

NK-92 cell line was acquired from the American Type Culture Collection and maintained in X-VIVO 10 (Lonza) supplemented with 20% FBS and 0.02 mM folic acid (Millipore Sigma) dissolved in 1 N NaOH, 0.2 mM myo-inositol (Millipore Sigma), and IL-2 (100 U/ml; PeproTech).NK cells were collected and after centrifugation at room temperature for 3 min at 300×g, pellets were resuspended in modified DM supplemented with 0.02 mM folic acid, 0.2 mM myo-inositol, and IL-2 (100 U/ml). Resuspended cells were introduced into the endothelial vessel and co-culture for 24h in excess medium.

### Neutrophil isolation

All blood samples were drawn according to institutional review board-approved protocols per the Declaration of Helsinki at the University of Wisconsin–Madison. Neutrophils were purified from whole blood using the MACSxpress Neutrophil Isolation Kit per the manufacturer’s instructions (Miltenyi Biotec) and residual red blood cells were lysed using MACSxpress Erythrocyte Depletion Kit (Miltenyi Biotec). Blood was drawn from a total of three nondiseased donors with informed consent obtained at the time of the blood draw according to requirements of the IRB. Prior to loading, the purified neutrophils were stained with calcein AM at 10 nM (Thermo Fisher) according to the manufacturer’s instructions. Neutrophils were resuspended in 50:50% v/v EMEM and EGM-2MV culture medium for co-culture with Caco-2 epithelium or resuspended in modified DM for co-culture with primary intestinal epithelium.

### Parasite cell culture and infection

*T. gondii* parasites were harvested and passaged using previously established protocols [113]. Briefly, *T. gondii* ME49 tachyzoites were propagated in human foreskin fibroblast (HFF) monolayers grown in Dulbecco’s modified Eagle medium (DMEM) containing 10% FBS and 1% Pen/Strep. Tachyzoites were harvested and pelleted by centrifugation at 2,200g for 10 min and resuspended in pre-warmed modified DM or 50:50% v/v EMEM and EGM-2MV culture medium prior to injection into the lumen of the epithelial tubes for infect studies.

### Immunofluorescence staining

For immunostaining, cells were fixed with 4% (v/v) paraformaldehyde (Alfa Aesar) for 20 min and permeabilized with 0.2% (v/v) Triton-X 100 (MP Biomedicals) for 10 min with three 1X PBS wash step between each solution. To reduce nonspecific background fluorescence from collagen, cells were incubated in 0.1 M glycine (Fisher Scientific, Pittsburgh, PA) for 30 min and washed with 1X PBS three times. Subsequently, cells were blocked with buffer solution (3% wt/v BSA and 0.1% v/v Tween 20 (Fisher Scientific)) overnight at 4 °C. Primary antibodies, **Table S2**, diluted in buffer solution were added to the microdevices and incubated at 4 °C for 2 days and washed with 1X PBS three times. Secondary antibodies, **Table S2**, were added to the buffer solution and incubated for 1 day at 4 °C. For cytoskeletal actin and nuclear staining, Alexa conjugated phalloidin (Thermo Fisher) and Hoechst 33342 (Thermo Fisher) at 1:100 were also added to the secondary antibody buffer solution. Lumens were then rinsed with 1X PBS three times over a two-day period.

### Image acquisition

Bright-field and fluorescent images were captured in a Nikon Ti Eclipse with a top stage incubator equipped with temperature and CO2 control (set at 37°C and 5%, respectively). Neutrophil kinetic parameters including end-to-end displacement and migration speed were analyzed with Fiji (https://imagej.net/Fiji) using the track-mate module. Confocal images were acquired at University of Wisconsin-Madison Optical Imaging Core using a Leica SP8 microscope.

### Optical metabolic imaging

A custom built inverted multiphoton microscope (Bruker Fluorescence Microscopy, Middleton, WI), was used to acquire fluorescence intensity and lifetime images. The equipment consists of an ultrafast laser (Spectra Physics, Insight DSDual), an inverted microscope (Nikon, Eclipse Ti), and a 40× water immersion (1.15NA, Nikon) objective. FAD fluorescence was isolated using an emission bandpass filter of 550/100 nm and excitation wavelength of 890 nm. NAD(P)H fluorescence was isolated using an emission bandpass filter of 440/80 nm and an excitation wavelength of 750 nm. The optical redox ratio was determined from the NAD(P)H and FAD lifetime data by integrating the photons detected at each pixel in the image to calculate the total intensity. For each pixel, the intensity of NAD(P)H was then divided by the intensity of FAD. Using Cell Profiler, an automated cell segmentation pipeline was created. This system identified pixels belonging to nuclear regions by using a customized threshold code. Cells were recognized by propagating out from the nuclei within the image. To refine the propagation and to prevent it from continuing into background pixels, an Otsu Global threshold was used. The cell cytoplasm was defined as the cell borders minus the nucleus. Values for NAD(P)H intensity, FAD intensity, and the optical redox ratio (NAD(P)H/FAD intensity) were averaged for all pixels within each cell cytoplasm. At least 100 cells per sample were analyzed, and every experiment was repeated at least three times.

### Cell retrieval from microdevice

To quantify gene expression related to proliferation, differentiation and a function of the intestinal epithelium primary intestinal epithelial cells were selectively retrieved from the intestinal tissue MPS consisting of epithelial and endothelial tubes. The upper half of the microdevice was removed to expose the collagen hydrogel. The hydrogel was then transferred to an Eppendorf tube containing 300 μl of type I collagenase (6 mg/ml). The sample was incubated on ice for 2 min to degrade the hydrogel and release the cells. Two microliters of biotinylated anti-EpCAM (Thermo Fisher Scientific) was added, and the sample was incubated at 4°C for 15 min. Ten microliters of SeraMAGS beads coupled to streptavidin was added, and the sample was incubated for another 10 min at 4°C. The SeraMAGS beads, with the epithelial cells (EpCAM-positive), were isolated using a magnet and lysed for PCR analysis. To quantify gene expression related to protective immunity, neutrophils were selectively retrieved from the intestinal tissue MPS 6h after introduction into the endothelial vessel. Non-adherent neutrophils were first collected from the endothelial vessels through the pippet-accesible ports. To isolate neutrophils that have migrated into the ECM gel, a 2-mm-diameter biopsy punch (Fisher Scientific) was used to cut out the hydrogel next to the endothelial vessel. The hydrogel punches were transferred to an Eppendorf tube and processed to separate EpCAM-positive epithelial cells, following the same protocol described above, leaving behind neutrophils in suspension. These neutrophils were mixed with neutrophils collected from the endothelial vessel and lysed for PCR analysis.

### Reverse transcription quantitative polymerase chain reaction

To study how primary intestinal epithelial cells and neutrophils adapted to the microenvironment within the intestinal tissue MPS, the expression of multiple genes related to different pathways was analyzed by RT-qPCR. Briefly, mRNA was isolated from cells using the Dynabeads mRNA DIRECT Purification Kit (Invitrogen). Isolated mRNA was quantified using a Qubit fluorometer (Thermo Fisher Scientific) and the Qubit RNA BR Assay Kit (Thermo Fisher Scientific). cDNA was produced using the High Capacity RNA-to-cDNA Kit (Applied Biosystem) and the cDNA was pre-amplified with SsoAdvanced PreAmp Supermix (Bio-Rad) using primers from Integrated DNA Technologies or Thermo Fisher Scientific, **Table S3-4**. cDNA was analyzed by RT-qPCR using iTaq Universal SYBR Green Supermix (Bio-Rad) or Roche Lightcycler master mix according to manufacturer’s protocols in Roche’s Lightcycler 480 II. Gene expression was normalized using the delta-delta Ct method. To quantify genomic DNA of T.gondii, infected epithelial tubes were digested as descrived above, and genomic DNA was extracted using TRIzol according to manufactures instructions. DNA was purified by phenol/chloroform extraction followed by ethanol precipitation. Genomic DNA was used as the template for preamplification and RT-PCR, as described above, using *T. gondii* primers (Table S2).

### Multiplex cytokine/chemokine assays and analysis

To measure NK cell-mediated cytokines secretion, media were collected from intestinal tissue MPS with *T. gondii* infected epithelium in the presence and absence of NK cells. Media were collected after 24h in culture and cytokine/chemokine concentrations analyzed using the Inflammation 20-Plex Human ProcartaPlex™ panel (EPX200-12185-901, Thermo Fisher) following the manufacturer’s guidelines. Data were collected on MAGPIX Luminex Xmap system (Luminex Corporation) using Luminex xPonent software. Concentration of each analyte was determined from a standard curve, generated by fitting a five-parameter logistic regression of median fluorescence on known concentrations of each analyte.

### Statistical analysis

Data were analyzed (Prism 9.0; GraphPad Software). The normal distribution assumption for statistical tests was confirmed by the Shapiro Wilk test. Statistical significance was assessed using Student’s t tests when comparing two conditions/groups and when comparing more than two groups, significance was assessed using one-way analysis of variance (ANOVA) corrected using the Tukey’s test. For nonparametric comparisons, a Mann-Whitney U test or a Kruskal-Wallis test was performed.

## Supporting information

Genes associated with normal proliferation and differentiation, or functional human intestinal epithelial cells were compared, Fig. S1-2

## Acknowledgments

We thank Dr. Evie Carchman from the Department of Surgery and UWCCC Biobank at the UW-Madison for the supply of small intestinal tissue, Huttenlocher Lab from the Department of Medical Microbiology and Immunology at the University of Wisconsin-Madison for facilitating blood draws from healthy donors. We also thank the UW Optical Imaging Core for the support on the 3D confocal imaging.

## Author contributions

M.H, J.A, K.P, M.S, S.K, L.J.K and D.J.B conceptualized and designed the research. M.H performed experiments and did data analysis with assistance from J.A, K.P and B.M.D. M.H, J.A. S.K, L.J.K and D.J.B wrote the manuscript. All authors reviewed the manuscript.

## Data and materials availability

All data needed to evaluate the conclusions in the paper are present in the paper and/or the Supplementary Materials. Additional data related to this paper may be requested from the authors.

