## Supplementary material for "Innate immune cell response to host-parasite interaction in a human intestinal tissue microphysiological system": Genes associated with normal proliferation and differentiation, or functional human intestinal epithelial cells were compared, Fig. S1-2

Figure S1.

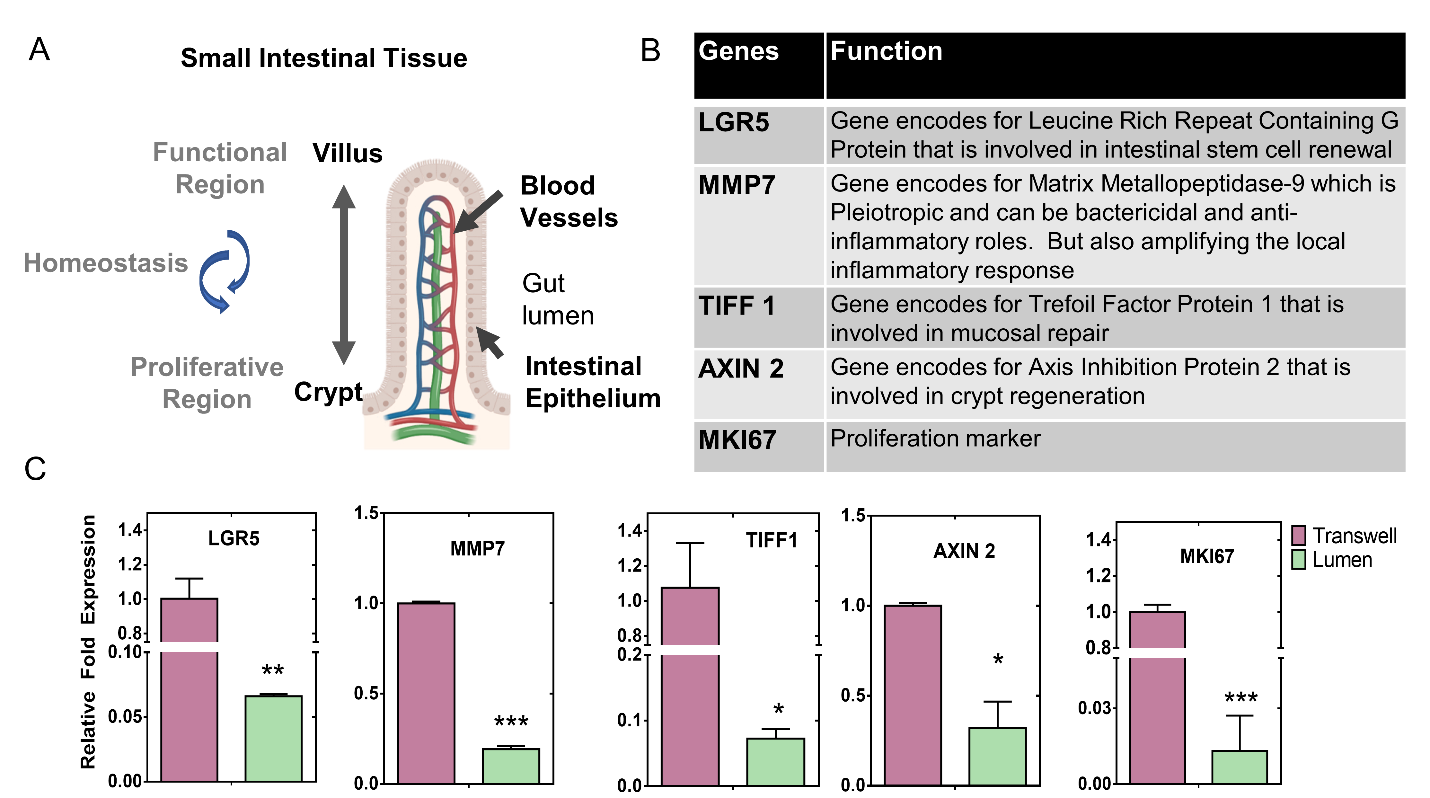

Fig. S1 Upregulated genes in primary intestinal epithelium cultured and differentiated in the human intestinal tissue MPS relative to standard culture methods

(A) Schematic representation showing the crypt-villus axis of the intestinal epithelium and the spatial distribution of proliferative and functional region. (B) Markers associated with proliferation and differentiation, or functional epithelium of the intestinal tissue. (C) Differential gene expression of the intestinal epithelium in the intestinal tissue MPS versus in transwell culture. Bar graph showing gene expression of downregulated genes, which include leucine-rich-repeat-containing G-protein-coupled receptor 5 (LGR5) for intestinal stem cells; Matrix Metallopeptidase-7 (MMP7) bactericidal and anti-inflammatory effects; Trefoil Factor 1 (TIFF1) for mucosal repair; Axis Inhibition Protein 2 (AXIN2) for crypt regeneration and proliferation marker (MKI67) for stem cells. Values are presented as mean ±SD from 4 independent experiments involving tubular or monolayer epithelium generated from human intestinal organoids where significance is expressed as ***p ≤ 0.001, **p ≤ 0.01 and *p ≤ 0.05).

Figure S2.

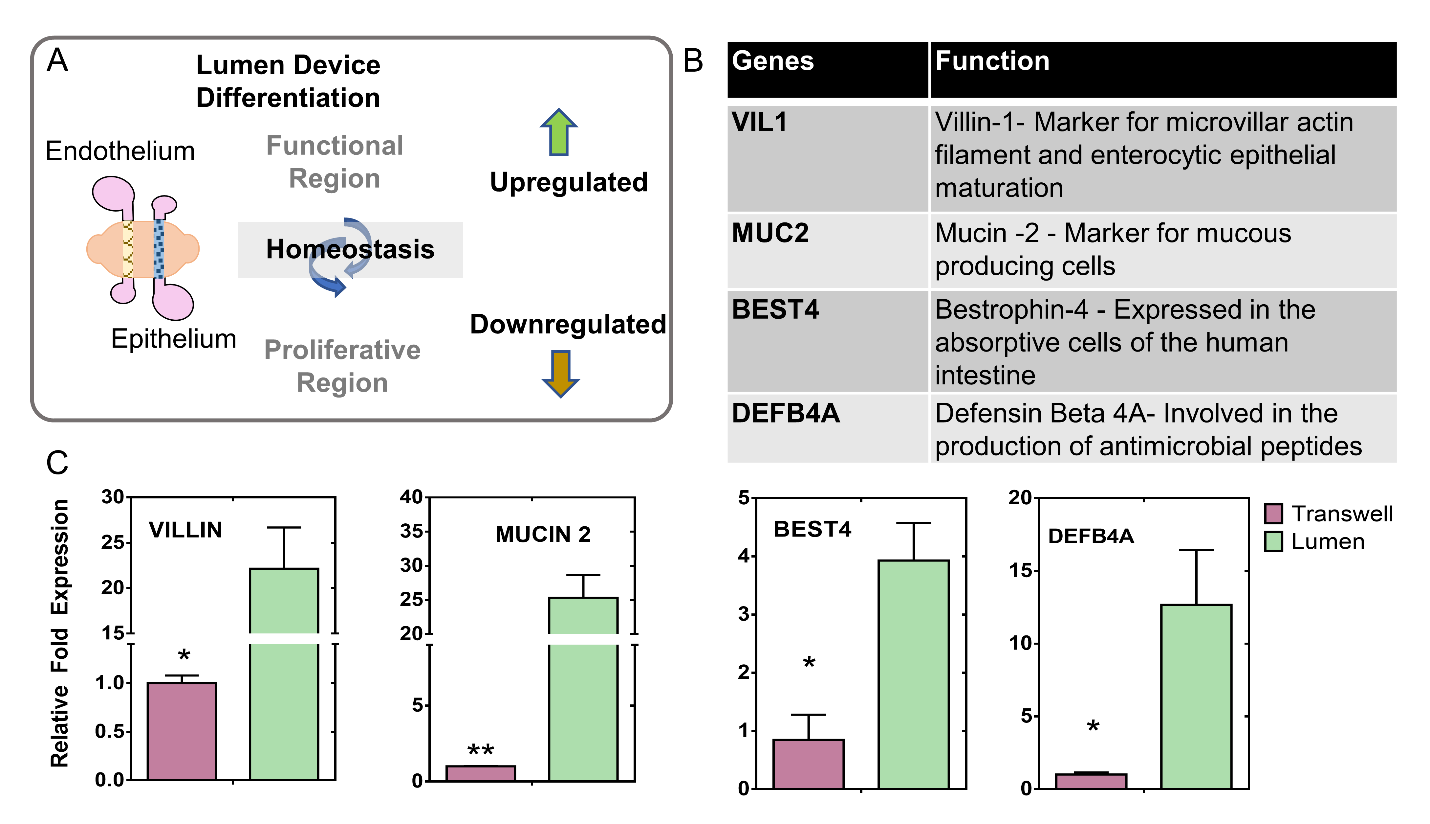

Fig. S2 Downregulated genes in primary intestinal epithelium cultured and differentiated in the human intestinal tissue MPS relative to standard culture methods

(A) Schematic representation showing upregulation and downregulation of genes in the epithelial tubes formed in the intestinal tissue MPS and their association with the proliferative and functional region of the intestinal epithelium. (B) Markers associated with proliferation and differentiation, or functional intestinal epithelium in the intestinal tissue. (C) Differential gene expression of the intestinal epithelium in the intestinal tissue model versus in transwell culture. Bar graph showing gene expression of upregulated genes, which include villin-1 (VIL1) for microvilli formation; mucin 2 (MUC2) for goblet cells; Bestrophin-4 (BEST4) for absorptive cells and β-defensin-4 (DEFB4) for antimicrobial peptides. Values are presented as mean ±SD from 4 independent experiments involving tubular or monolayer epithelium generated from human intestinal organoids where significance is expressed as **p ≤ 0.01 and *p ≤ 0.05).

Figure S3.

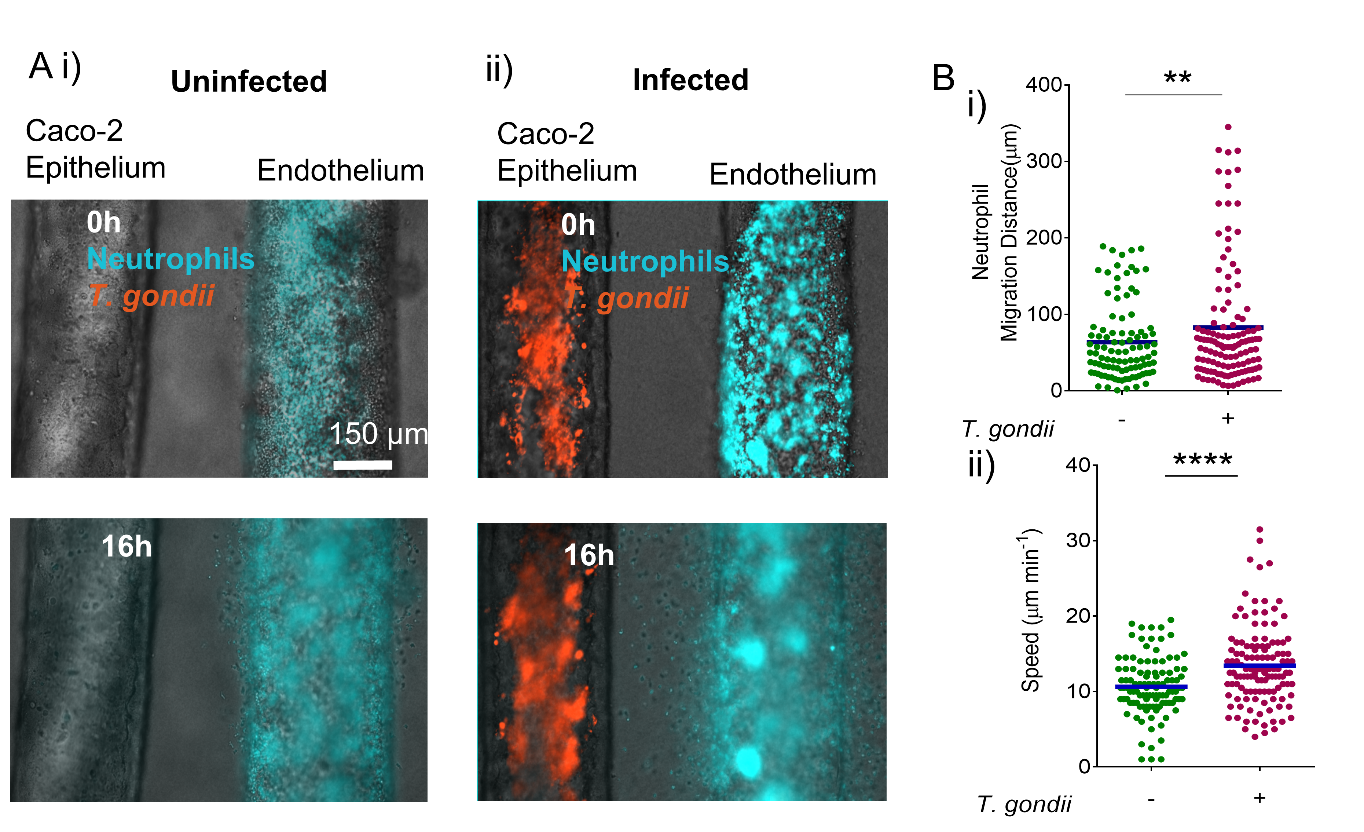

Fig. S3 Dynamic analysis of neutrophils extravasation and trafficking towards *T.gondii* infected epithelium

**(A)** Combined brightfield and immunofluorescence images showing the interface between the gut epithelium and endothelium in the intestinal tissue MPS. Neutrophils were introduced into the endothelial vessel and monitored for their extravasation and migration behavior over a 16-h period (i). Caco-2 epithelial tubes were infected with m-cherry tagged ME49 *T. gondii* for 48h before introducing neutrophils into the endothelial vessel (ii). Neutrophils were seen to extravasate and migrate towards the infected epithelium. **(B)** Grouped scatter plot showing migration distance (i), as measured by end-to-end displacement following extravasation of neutrophils, and the average speed towards a *T.gondii*-infected epithelium. Each dot represents the migration distance and speed of a single neutrophil from the endothelial vessel boundary, significance is expressed as ****p ≤ 0.0001 and **p ≤ 0.01.

Figure S4.

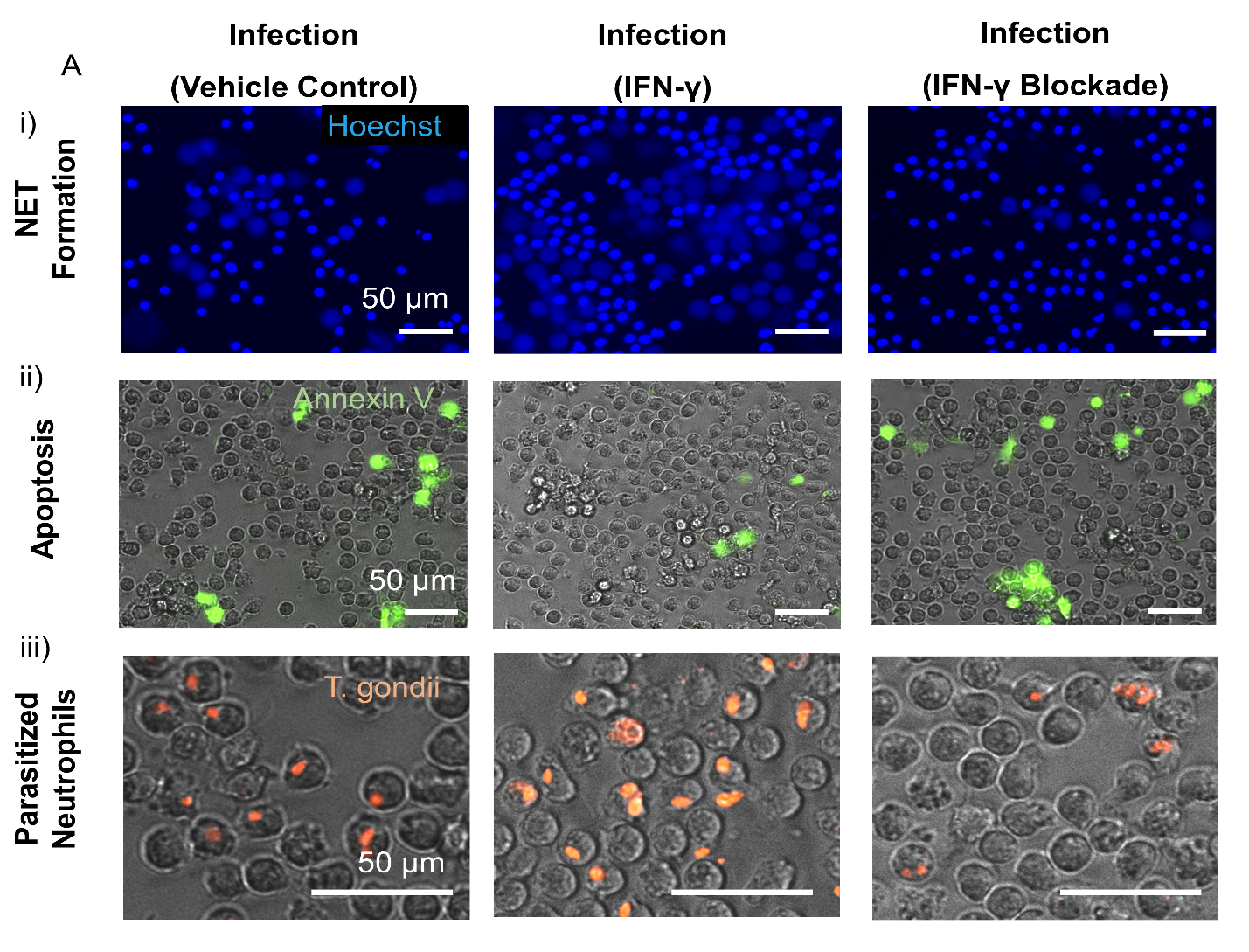

Fig. S4 Influence of IFN- γ stimulation and inhibition on neutrophil function at the site of *T. gondii* infected epithelium (A) Representative fluorescence images showing NET formation, apoptosis and parasitization of neutrophils from interacting with *T.gondii-*infected epithelium in response to IFN-γ stimulation and inhibition. Caco-2 epithelial tubes were infected with m-cherry tagged ME49 *T. gondii* for 72h before introducing neutrophils into the lumen of the tube. Neutrophils were co-cultured with the infected epithelium for 6 h prior to collection in well-plate and imaging. Fluorescences image showing neutrophils stained with Hoechst 33342 for DNA, decondensed nuclei indicating NET formation (i). Combined brightfield and fluorescence image showing neutrophils stained with Apopxin Green (abcam) as indicator of cells undergoing apoptosis, (ii). Combined brightfield and fluorescence image showing parasitized neutrophils where direct interaction with *T. gondii* parasite is observed (iii).

Figure S5.

Fig.S5 Optical redox ratio assessment in IFN-γ primed neutrophils during T. gondii infection.

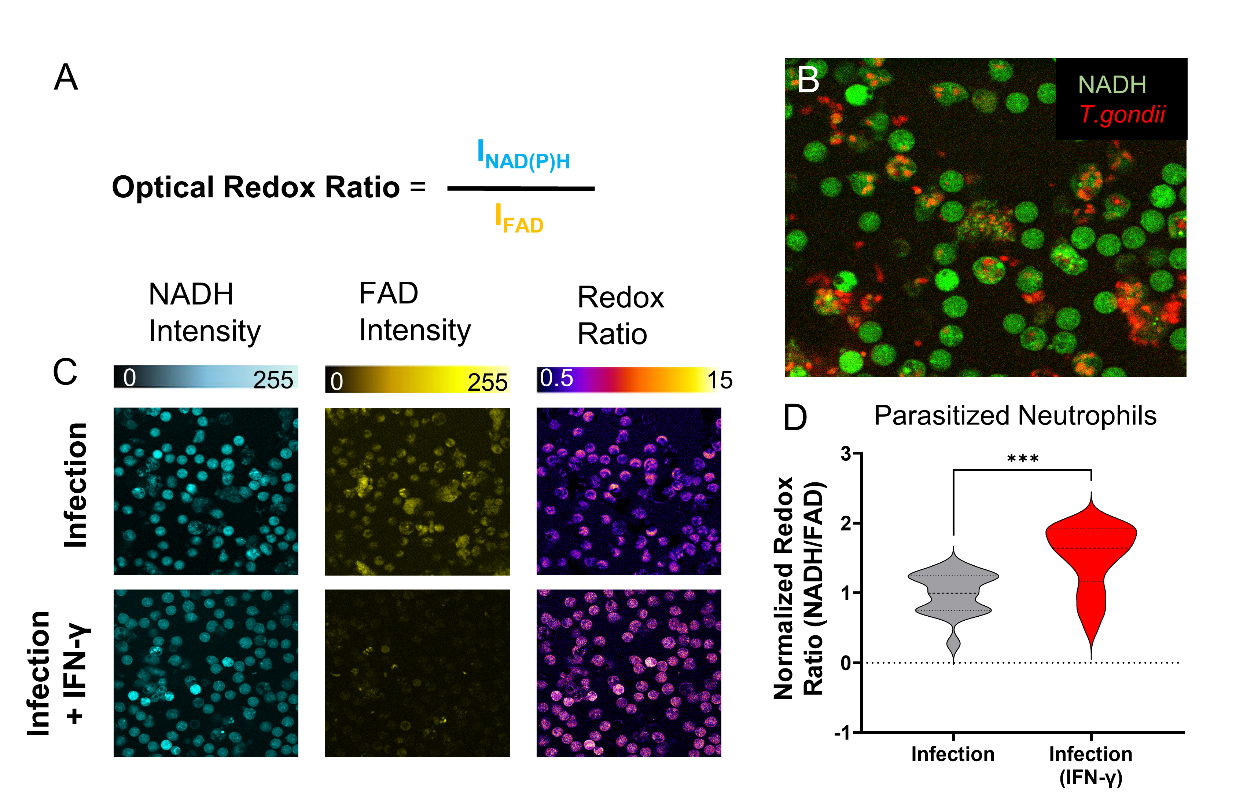

**Optical redox ratio assessment in IFN-γ primed neutrophils during *T. gondii* infection. (A)** The optical redox ratio, defined as the autofluorescence intensity of NADH divided by that of FAD, quantifies relative rates of cellular glycolysis and oxidative phosphorylation. Optical metabolic imaging was used to visualize intracellular NAD(P)H and FAD fluorescence intensities of neutrophils in infected systems where the neutrophils were directly introduced into the epithelium to increased instances of direct-contact interaction with *T. gondii* in the lumen of the infected epithelium. **(B)** Representative images showing autofluorescence intensity of NADH (green) in neutrophils from in a *T.gondii* (red) -infected epithelium in the intestinal tissue MPS. A number of parasitized neutrophils can be observed, as indicated by white arrows. **(C)** Representative images showing NADH, FAD and the Redox ratio intensities in neutrophils from intestinal tissue MPSs without and with IFN-γ stimulation. **(D)** Violin plots showing the analysis of neutrophil redox ratio in parasitized cells based on NAD(P)H and FAD intensity (asterisk denotes P value of ≤ 0.05).

Figure S6.

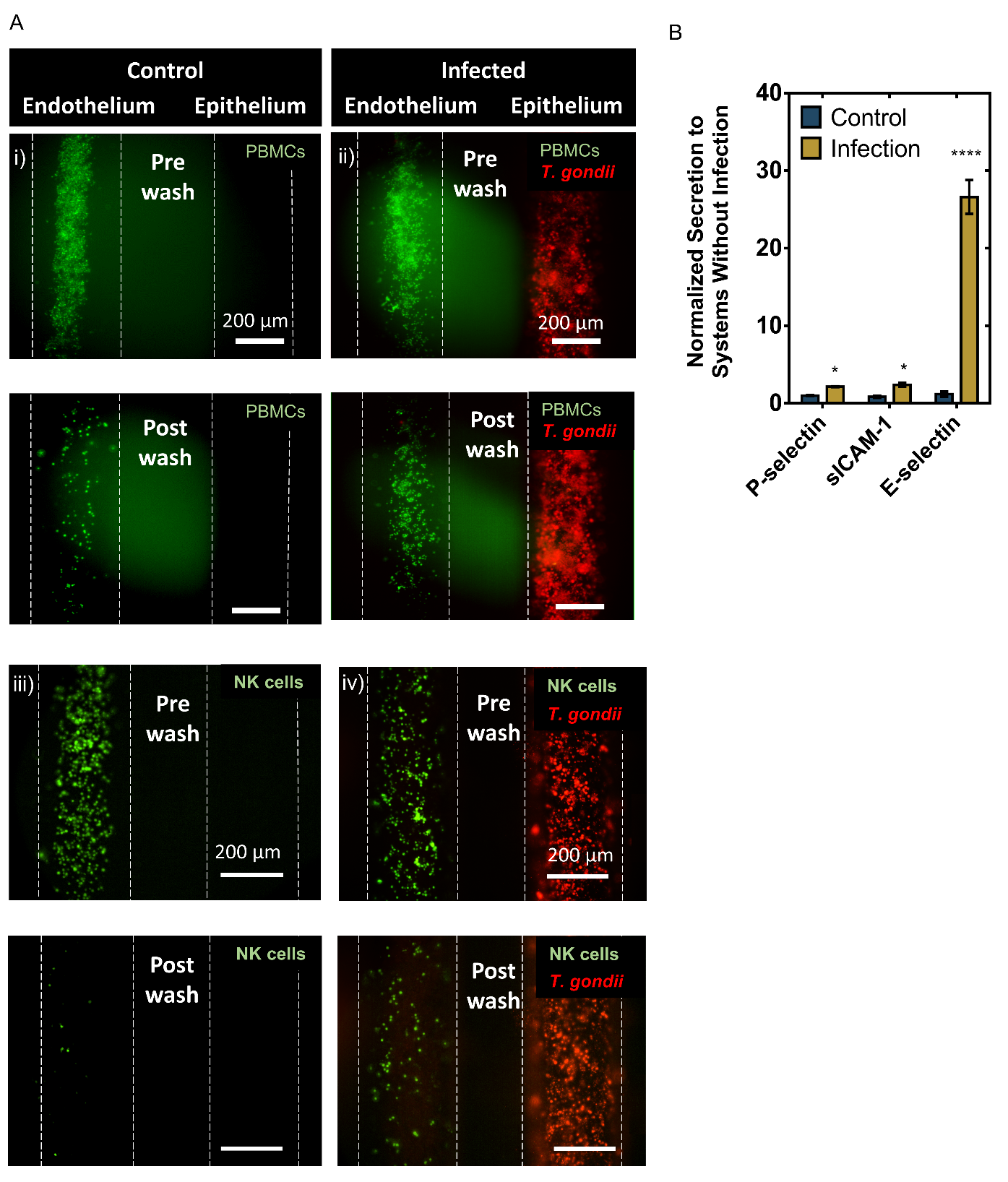

Fig. S6 Effects of epithelial infection by *T. gondii* on immune cell adhesion to endothelium. (A) PBMCs were added to the endothelium 48 hpi with ME49 *T. gondii* and washed away after 2h to evaluate endothelial adhesion of immune cells. Fluorescence image showing differences in primary PBMC adhesion events (pre wash and post wash) to the endothelial vessel in response to control (i) and *T.gondii*-infected (ii), Caco-2 epithelium. Similarly, NK cell adhesion was also examined. Fluorescence image showing differences in NK 92 cell adhesion events (pre wash and post wash) to the endothelial vessel in response to control (iii) and *T.gondii*-infected (iv), Caco-2 epithelium. (B) Increased levels of soluble adhesion molecules, sP-selectin, sICAM-1, and sE-selectin were also found in the culture media of infected systems. Cytokine concentrations measured in media collected from the intestinal tissue MPS consisting of endothelial vessel and primary intestinal epithelial tubes infected with *T. gondii* normalized to control, uninfected systems*.* In all cytokine measurement experiments nine devices were prepared on two different days for each paired conditions (infected vs. control, and infection in the absence vs. presence of NK cells), media from three devices were pooled to make one replicate. Significance is expressed as ****p ≤ 0.0001 and *p ≤ 0.05).

**Table S1. Top 10 most relevant GO terms (Biological Processes) associated with genes analyzed in neutrophils with their corresponding FDR-adjusted P value**

| **Biological Processes^a^** | **FDR-adjusted P** |
| --- | --- |
| Cytokine Production | 9.84E-23 |
| Positive Regulation Of Cytokine Production | 6.21E-22 |
| Regulation Of Leukocyte Mediated Immunity | 4.13E-21 |
| Leukocyte Differentiation | 4.17E-21 |
| Positive Regulation Of Immune Effector Process | 4.40E-21 |
| Response To Cytokine | 6.05E-21 |
| Defense Response | 1.01E-20 |
| Immune Effector Process | 1.84E-20 |
| Adaptive Immune Response | 8.99E-20 |
| Regulation Of Immune Effector Process | 8.99E-20 |

^a^ Based on the GSEA molecular signatures database

**Table S2. Primary and secondary antibodies used for immunofluorescent staining**

| Primary antibody | Company code | Dilution | **Secondary**  **antibody** | **Dilution** |
| --- | --- | --- | --- | --- |
| CD31 | Abcam AB28364 | 1:20 | ^a^Alexa Fluor 488 anti-rabbit (Thermo Fisher Scientific, A-11008) | 1:100 |
| MUC2 | Santa Cruz Biotechnology, sc-515032 | 1:50 | ^a^Alexa Fluor 488 anti-mouse (Thermo Fisher Scientific, A-11001) | 1:100 |
| Villin | Novus Biologicals, NBP2-53201 | 1:100 | ^a^Alexa Fluor 647 anti-mouse (Thermo Fisher Scientific, A-21235) | 1:100 |
| ZO-1 | Thermo Fisher Scientific, 61-7300 | 1:25 | ^a^Alexa Fluor 568 anti-rabbit (Thermo Fisher Scientific, A-11011) | 1:100 |
| E-cadherin | BD Biosciences, 610182 | 1:25 | ^a^Alexa Fluor 488 anti-mouse (Thermo Fisher Scientific, A-11001) | 1:100 |
| FABP1 | Sigma Aldrich, HPA028275 | 1:25 | ^a^Alexa Fluor 568 anti-rabbit (Thermo Fisher Scientific, A-11011) | 1:100 |

^a^ Goat source

Table S3. Primer set sources for gene expression analysis of primary intestinal epithelium culture in intestinal tissue MPS and in transwells

| **Gene** | | **Source** |
| --- | --- | --- |
| Genes associated with proliferation and differentiation, or functional intestinal epithelial cells. | MKI67 | Thermo Fisher Scientific, Hs04260396_g1 |
|  | Axis inhibition protein 2 (Axin2) | Thermo Fisher Scientific, Hs00610344_m1 |
|  | Trefoil factor 1 (TFF1) | Thermo Fisher Scientific, Hs00907239_m1 |
|  | Sucrase-isomaltase (SI) | Thermo Fisher Scientific, Hs00356112_m1 |
|  | Villin | Thermo Fisher Scientific, Hs01031739_m1 |
|  | Mucin 2 (MUC2) | Thermo Fisher Scientific, Hs03005103_g1 |
|  | Hypoxia-inducible factor (HIF-1α) | Thermo Fisher Scientific, Hs00153153_m1 |
|  | Leucine-rich-repeat-containing G-protein-coupled receptor 5 (LGR5) | Thermo Fisher Scientific, Hs00969422_m1 |
|  | Matrix Metallopeptidase-7 (MMP7) | Thermo Fisher Scientific, [Hs01042796_m1](https://www.thermofisher.com/taqman-gene-expression/product/Hs01042796_m1?CID=&ICID=&subtype=) |
|  | Lysozyme (LYZ) | Thermo Fisher Scientific, Hs00426232_m1 |
|  | Tight junction protein-1 (TJP1) | Thermo Fisher Scientific, Hs01551861_m1 |
|  | β-defensin-1 (DEFB1) | Thermo Fisher Scientific, [Hs00174765_m1](https://www.thermofisher.com/taqman-gene-expression/product/Hs00174765_m1?CID=&ICID=&subtype=) |
|  | β-defensin-4-alpha (DEFB4) | Thermo Fisher Scientific, [Hs00823638_m1](https://www.thermofisher.com/taqman-gene-expression/product/Hs00823638_m1?CID=&ICID=&subtype=) |
|  | Bestrophin-4 (BEST4) | Thermo Fisher Scientific,  [Hs00396114_m1](https://www.thermofisher.com/taqman-gene-expression/product/Hs00396114_m1?CID=&ICID=&subtype=) |
| Housekeeping  (Reference genes) | GAPDH | Thermo Fisher Scientific, Hs01922876_m1 |
|  | HPRT | Thermo Fisher Scientific, Hs02800695_m1 |
|  | RPLP0 | Thermo Fisher Scientific, Hs99999902_m1 |

**Table S4. Primer sequences for neutrophil gene expression analysis**

| **Gene** | | **Source** |
| --- | --- | --- |
| Genes associated protective immunity. | Interleukin – 10 (IL10) | Integrated DNA Technologies,  Hs.PT.58.15400284 |
|  | MYD88 innate immune signal transduction adaptor | Integrated DNA Technologies,  Hs.PT.58.40601199.gs |
|  | Tumor necrosis factor (TNF) | Integrated DNA Technologies,  Hs.PT.58.45380900 |
|  | Interleukin – 1beta (IL1B) | Integrated DNA Technologies,  Hs.PT.58.1518186 |
|  | Interleukin – 6 (IL6) | Integrated DNA Technologies,  Hs.PT.58.40226675 |
|  | Interleukin – 1alpha (IL1A) | Integrated DNA Technologies,  Hs.PT.58.40913627 |
|  | Nitric oxide synthase 2 (NOS2) | Integrated DNA Technologies,  Hs.PT.58.45668131 |
|  | Interleukin – 12A (IL12A) | Integrated DNA Technologies,  Hs.PT.58.1687020 |
|  | [NLR family pyrin domain containing 3](https://www.ncbi.nlm.nih.gov/gene/114548)  (NLRP3) | Integrated DNA Technologies,  Hs.PT.58.39497108 |
|  | Interferon gamma (IFNG) | Integrated DNA Technologies,  Hs.PT.58.24522521 |
|  | Toll-like receptor-2 (TLR2) | Integrated DNA Technologies,  Hs.PT.58.26767404 |
|  | Toll-like receptor-4 (TLR4) | Integrated DNA Technologies,  Hs.PT.58.38700156.g |
|  | Transforming growth factor beta 1 (TGFB1) | Integrated DNA Technologies,  Hs.PT.58.39813975 |
|  | Interleukin – 4 receptor (IL4R) | Integrated DNA Technologies,  Hs.PT.58.23069040 |
|  | Interleukin – 2 (IL2) | Integrated DNA Technologies,  Hs.PT.58.1142676 |
|  | Interferon alpha 1 (IFNA1) | Integrated DNA Technologies,  Hs.PT.58.46311748.g |
|  | Interferon beta 1 (IFNB1) | Integrated DNA Technologies,  Hs.PT.58.39481063.g |
| *T.gondii* genes | Housekeeping gene: alpha-tubulin (TUB1A) | Integrated DNA Technologies,  Forward: 5′ –GACGACGCCTTCAACACCTTCTTT– 3′ Reverse: 5′ –AGTTGTTCGCAGCATCCTCTTTCC– 3′ |
|  | SAG1 | Integrated DNA Technologies,  Forward: 5′ –TGCCCAGCGGGTACTACAAG– 3′ Reverse: 5′ –TGCCGTGTCGAGACTAGCAG– 3′ |
| Housekeeping for mammalian cells  (Reference genes) | GAPDH | Integrated DNA Technologies,  Hs.PT.39a.22214836 |
|  | HPRT | Integrated DNA Technologies,  Hs.PT.39a.22214821 |
|  | RPLP0 | Integrated DNA Technologies,  Hs.PT.39a.22214824 |

Movie S1.

**Neutrophil extravasation and trafficking towards control intestinal epithelium**

Movie S2.

**Neutrophil extravasation and trafficking towards**
